# Targeted rescue of synaptic plasticity improves cognitive decline after severe systemic inflammation

**DOI:** 10.1101/2021.03.04.433352

**Authors:** Benedikt Grünewald, Jonathan Wickel, Nina Hahn, Franziska Hörhold, Hanna Rupp, Ha-Yeun Chung, Holger Haselmann, Anja S. Strauss, Lars Schmidl, Nina Hempel, Lena Grünewald, Anja Urbach, Michael Bauer, Klaus V. Toyka, Markus Blaess, Ralf A. Claus, Rainer König, Christian Geis

## Abstract

Sepsis-associated encephalopathy (SAE) is a frequent complication in patients with severe systemic infection resulting in acute brain dysfunction and incapacitating long-term sequelae. SAE includes delirium, premature death, post-traumatic stress disorder, and major long-term cognitive impairment. The underlying pathophysiology of SAE is largely unresolved and specific treatment options are missing. We induced the peritoneal contamination and infection (PCI) sepsis model in 769 mice and compared these with 259 control mice. We found that experimental sepsis causes synaptic pathology in the brain characterized by severely disordered synaptic plasticity with reduced long-term potentiation, changes in CA1 pyramidal neuron dendritic spines, and behavioral abnormalities indicating cognitive dysfunction. Using electrophysiology, we found reduced frequency of quantal and spontaneous excitatory postsynaptic currents whereas amplitudes of miniature, spontaneous, and evoked excitatory currents were increased, pointing towards a homeostatic synaptic scaling mechanism. Corresponding to dysfunctional excitatory synaptic function we discovered downregulation of genes related to neuronal and synaptic signaling in the brain, including the gene for activity-regulated cytoskeleton-associated protein (*Arc/Arg3.1*), members of the transcription-regulatory EGR gene family, and the gene for dual-specificity phosphatase 6 (*Dusp6*). At the protein level, ARC expression and MAP kinase signaling in the brain were affected. For targeted rescue of dysfunctional synaptic signaling and plasticity, we overexpressed ARC in the hippocampus by microinjection of an adeno-associated virus containing a neuron-specific plasmid of the ARC transgene. Hereby we achieved recovery of defective synaptic plasticity in the hippocampal Schaffer collateral-CA1 pathway and improvement of memory dysfunction. Using a different rescue paradigm, PCI mice were subjected to enriched environment providing multiple activating stimuli. Enriched environment led to restoration of disordered long-term potentiation and memory, thus demonstrating the potential for activity-induced improvement. Together, we identified synaptic pathomechanisms of SAE after severe systemic infection and provide a conceptual approach to treat SAE-related disease mechanisms which may be applicable to patients afflicted with SAE.

## Introduction

Growing evidence suggests that severe systemic infections may lead to long-term cognitive dysfunction with the risk of developing dementia.^1,2^ Sepsis as the most severe manifestation of systemic infection is defined as a life-threatening organ dysfunction caused by an overwhelming and dysregulated host response to inflammation.^3^ Sepsis is a world-wide health problem with increasing incidence and high mortality. Recent epidemiological studies revealed an incidence rate of 437 hospital-treated sepsis patients per 100.000 person-years and an estimated global incidence of 31.5 million sepsis cases with 5.3 million deaths annually.^4^ Sepsis-associated encephalopathy (SAE) is a frequent and severe complication and often presents as the first and leading disease sign. SAE occurs in an estimated 70% of patients with severe sepsis and has been identified as an independent risk factor for increased mortality.^5–7^ Long-term sequelae include marked cognitive deficits comparable to those found in patients with beginning Alzheimeŕs dementia.^8^

The mechanisms of SAE are likely multifactorial. Postmortem findings in patients and in animal models provided evidence for severe systemic inflammation, BBB disruption,^9,10^ and activation of intracerebral immune cells, e.g. microglia and hematogenous macrophages.^11–14^ This inflammatory milieu in the brain leads to the release of cytokines,^15,16^ chemokines,^17^ oxygen radicals^18^ and reactive nitrogen intermediates.^19^ In addition, ischemia,^20^ hemorrhages,^21^ abnormal alterations in cerebral homeostasis, and dysregulation of cholinergic neurotransmission^22^ may all contribute to the development of SAE (reviewed in Chung ^23^, Gofton^7^ and Mazeraud.^24^). Yet, the pathophysiology underlying SAE and cognitive decline after systemic infection in humans is only incompletely understood. Specifically, it is unclear how severe systemic inflammation may mediate persistent neuronal and synaptic dysfunction finally leading to neurocognitive defects. To date, no specific treatment options have been defined.

On a broader scale, neuropsychiatric defects are now increasingly recognized in other infectious disorders, e.g. systemic viral infection by SARS-CoV2. Indeed, neuropsychiatric sequelae including cognitive dysfunction, is a common and serious manifestation following COVID-19.^25^

Using a multidisciplinary approach, we here focused on the synaptic pathophysiology following severe systemic infection in a mouse model of peritoneal polymicrobial sepsis. We identified severely impaired synaptic pathways and disordered plasticity. Based on these findings we were able to establish therapeutic rescue strategies finally resulting in the amelioration of neuronal functionality and overall disease signs. Targeting defective regulatory mechanisms of synaptic plasticity offers conceptual advances for translational strategies in treating the sequelae of SAE.

## Materials and methods

### Mouse model of experimental sepsis

All animal experiments were approved by the state authorities of Thuringia (UKJ-02-085-14) and were performed in accordance with animal welfare and the ARRIVE guidelines.^26^ All efforts were made to minimize animal suffering and to keep the number of mice used as low as possible. Experimental sepsis was induced by the standardized and established peritoneal contamination and infection (PCI) mouse model as described previously.^27^ In brief, PCI mice received an intraperitoneal injection of human fecal slurry (diluted 1:4 in saline solution, 3 µl/g B.W.) with a 21-gauge cannula. Control mice were injected with 100µl of 0.9 M saline solution. We exclusively used male C57Bl6J mice as PCI disease severity may vary in female mice due to the menstrual cycle and hormonal fluctuations. Mice were closely monitored and the Clinical Severity Score (CSS, grade 1 to 4) was assessed as described previously^27^: grade 1: No signs of illness, active and curious, quick movements upon exogenous stimuli, normal posture; grade 2: Low-grade illness, less active with occasional interruptions in activity, reduced alertness, but adequate response to exogenous stimuli, posture slightly hunched; grade 3: moderately severe illness, slow and sleepy, moves with difficulty, limited and delayed reaction to exogenous stimuli, hunched posture; severe illness, mouse lethargic, motionless, no spontaneous movements, no reaction to exogenous stimuli, severely hunched posture. Mice were euthanized when CSS grade 4 was reached at two consecutive timepoints within 3 hours. To avoid premature death due to severe peritonitis, mice received the first subcutaneous (s.c.) injection of the beta-lactam antibiotic meropenem (650 µl, 1mg/ml) when the CSS reached a point value of 3. Subsequently, meropenem was injected s.c. twice daily for 7 days and once per day at the following three days to increase long-time survival after PCI. Thereafter, enrofloxacin (Baytril 2,5%, Bayer AG, Germany) was added to the drinking water with saccharose added in a final concentration of 2 mg/ml until the end of the experiment. Control animals also received equal amounts of the same antibiotics. For the experimental sepsis groups, only PCI animals with severe sepsis (cumulative 5 day-CSS of ≥ 11.5) were included in further experiments. A total of 1028 mice were used in the experiments (769 PCI, 259 control).

### Stereotactic intrahippocampal microinjection

Stereotactic surgery for injection of adeno-associated viruses (AAV) into the hippocampus was performed as previously described.^28^ Briefly, mice were deeply anesthetized with 1.5 to 2.0% of isoflurane/oxygen and the head was then fixed into a stereotactic apparatus (Lab StandardTM, Stoelting, Wood Dale, IL, USA). The skull was exposed by a midline incision, stereotactic injection sites were marked, and holes were drilled with a dentist driller (Foredom, Bethel, CT, USA). Injections of AAV were performed at three sites within the hippocampus: coordinates in mm with respect to bregma (AP/L/DV): 1st (2.8/2.8/2.8); 2nd (2.1/1.4/1.4); 3rd (2.1/1.4/2.1). Very thin injection pipettes (resistance > 20 MΩ) were pulled from borosilicate glass tubes (WPI, Sarasota, FL, USA, #4878) and filled with AAV preparation (concentration 1×10^10^ genome copies/µl) in 0.9 M saline solution. The injection pipette was placed at stereotactic coordinates and 1 µl of AAV containing solution was injected with an injection speed of 0.4 nl/s by a microprocessor controlled nanoliter injector (Nanoliter 2000 + SYS-Micro4 Controller, WPI, Sarasota, FL, USA) per injection site. Tissue damage in the target region was minimized by low injection volume and slow injection speed. After injections the skin of the animals was sutured and mice were taken back to their home cage.

### Behavioral analyses

Behavioral analyses included standardized test procedures, i.e. open field, elevated plus maze, Morris water maze, and Barnes maze. Tests were performed at 8 weeks after PCI when surviving mice had recovered from systemic disease signs and had reached a steady state of long-term behavioral dysfunction to be studied. Detailed procedures are described in Supplementary Material and Methods.

### Enriched environment

From day 10 after PCI or sham injection, mice were housed in an enriched environment (EE) cage (6 animals/cage, 85 cm × 75 cm × 40 cm; groups PCI EE and SHAM EE) with stimuli comprising objects (e.g. ladders and tunnels, running wheel, nesting material, and climbing boards). The configuration of the objects was changed on a weekly basis.^29^ The respective non-stimulated control groups (PCI NE and SHAM NE) remained in the normal standard housing conditions all over the experiment without additional stimulation of motor activity (2 animals/cage; 54 cm × 38 cm × 19 cm).

### Electrophysiology: patch-clamp and local field potential recordings

#### Preparation of acute hippocampal slices

For field potential (fEPSP) recordings acute coronal hippocampal slices were prepared from adult mice. Mice were sacrificed under isoflurane anesthesia by decapitation and brains were dissected and sliced (400µm) using a Leica VT 1200 microtome in ice-cold high-sucrose solution (in mM): 20 NaCl, 25 NaHCO_3_, 10 glucose, 150 sucrose, 4 KCl, 1.25 NaH_2_PO_4_, 0.5 CaCl_2_, 7 MgCl_2_. Slices were stored in artificial cerebrospinal fluid #1 solution (aCSF #1); in mM): 124 NaCl, 26 NaHCO_3_, 10 glucose, 3.4 KCl, 1.2 NaH_2_PO_4_, 2 CaCl_2_, 2 MgSO_4_ at 32°C for 30 min and subsequently at room temperature before transferring slices into the recording chamber.

For whole-cell patch-clamp recordings previously published protocols were adapted.^30,31^ Isoflurane-anesthetized mice were transcardially perfused with ice-cold protective aCSF #2 (in mM): 95 N-methyl-D-glucamine, 30 NaHCO_3_, 20 HEPES, 25 glucose, 2.5 KCl, 1.25 NaH_2_PO_4_, 2 thiourea, 5 Na-ascorbate, 10 MgSO_4_, 0.5 CaCl_2_ and 12 N-acetyl-L-cysteine. Brains were sliced according to previous protocols (‘magic cut’)^32^ and the slices were subsequently stored for 12 minutes in the same solution at 34°C. Afterwards slices were transferred to aCSF #3 solution (in mM): 125 NaCl, 25 NaHCO_3_, 25 glucose, 2.5 KCl, 1.25 NaH_2_PO_4_, 2 CaCl_2_, 1 MgCl_2_ supplemented with (in mM) 2 thiourea, 5 Na-ascorbate, 3 Na-pyruvate and 12 N-acetyl-L-cysteine. All solutions were continuously purged with carbogen (95% O_2_, 5% CO_2_) and the pH was maintained at 7.3.

#### Whole-cell patch clamp analyses in the dentate gyrus

Whole cell patch clamp recordings were performed using a EPC10 amplifier (HEKA, Germany) as described previously.^33^ During the experiment slices were superfused with aCSF#3 at RT. Dentate granule cell were visually identified and patched using glass pipettes (3 – 4 MΩ) filled with intracellular solution (in mM) 115 potassium gluconate, 40 KCl, 10 Hepes, 0.1 EGTA, 2 Na_2_ATP, 2 MgCl_2_. Excitatory postsynaptic currents (EPSCs) were recorded in the presence of bicucillin and CGP 52432. For the measurement of miniature EPSCs TTX (1 µM) was added. Minimal stimulation at 0.3 Hz applied via an aCSF #3-filled glass pipette within the medial perforant path was used to evoke eEPSCs. Stimulation of minimal evoked eEPSCs was determined by a synaptic failure rate of 50 % as reported previously.^34^

#### Analysis of long-term potentiation (LTP) by recording fEPSPs

Measurements of fEPSPs at the Schaffer collateral (SC) - CA1 synapses were performed as described.^35^ Slices were perfused with aCSF1 at 2.5 ml/min at 32-33°C. Stimulation within SC was performed using a stimulus isolation unit (Isoflex A.M.P.I, Jerusalem, Israel) and custom made bipolar platinum electrodes. Glass electrodes filled with aCSF#1 in the stratum radiatum were used to record the fEPSPs. Stimulation strength was varied between 25 and 400µA to record the input-output curve. For subsequent recordings the stimulation strength was adjusted by reducing the stimulation by 10% from the lowest strength which evoked a fEPSP with a discernable population spike. Stimuli were delivered at 0.03 Hz. LTP was induced after a baseline recording of 15 min by 20 theta bursts of 4 pulses of 100 Hz with an interval of 200 ms. After induction of LTP fEPSPs were recorded for 30 minutes. The fEPSP slopes were measured directly after the fiber volley to the peak or until the beginning of a clearly distinguishable population spike.

#### Analysis of LTP by whole cell patch clamp recording in CA1 pyramidal neurons

Whole-cell patch clamp recordings of CA1 pyramidal cells were performed in aCSF#2 with 100 µM picrotoxin. Glass pipettes were filled with internal solution (in mM): 115 potassium gluconate, 20 KCl, 10 HEPES, 4 Mg-ATP, 0.3. Na-GTP, 10 Na-phosphocreatine, 0.001 CaCl_2_, (pH 7.4, 280-290 mosmol/l). SC were stimulated using a glass pipette filled with aCSF#2 and a stimulus duration of 70 µs. Input-output curves were measured for every cell and stimulation strength was adjusted to evoke an EPSP of 30-50% of the maximum EPSP. After a baseline recording of 10 min at a frequency of 0.05 Hz, spike timing-dependent plasticity was induced by pairing a presynaptically evoked EPSP with 4 action potentials induced by postsynaptic current injection (3 ms duration, 1 nA, 200 Hz) according to previously published protocols.^36^. This paring was repeated 35 times at 0.5Hz. Afterwards EPSPs were recorded for further 30 min. EPSP slopes were calculated from the initial 2 ms after EPSP onset. For final analysis, the data were normalized to the average of the baseline recording^36^

#### Data acquisition and data analysis of electrophysiological recordings

All data were obtained using an EPC10 amplifier controlled by Patchmaster software (HEKA, Germany). mEPSCs were analyzed using Clampfit 10 (Molecular Devices, USA). All other data were analyzed using NeuroMatic plugin (http://www.neuromatic.thinkrandom.com) of Igor Pro Software (Wavemetrics, Portland, OR, USA).

### Generation of Arc expressing constructs and of negative control Adeno-associated viral (pAAV) constructs

To generate a pAAV construct that codes for the murine Arc mRNA (indicated as ARC-AVV), the Arc cDNA open reading frame (ORF) was cut from a pCMV6-Entry_Arc vector (Origene, MR206218) by restriction digest using EcoRI and XhoI (Fermentas) and inserted into a p-AAV-Syn-iCre-2A-Venus vector^37^ at the respective restriction sites using T4 Ligase (Fermentas). As negative control adaptor oligonucleotides have been generated by annealing two complementary noncoding sequences with EcoRI and XhoI restriction site overhangs (Neg(+): AATTgcatttgtactgacatct; Neg(-): TCGAagatgtcagtacaaatgc). This adaptor was ligated with the p-AAV-Syn-iCre-2A-Venus vector and linearized by EcoRI and XhoI treatment (indicated as CTRL AVV). Sequences of constructs were verified by Sanger sequencing at Eurofins Genomics, Ebersberg, Germany.

#### Virus production, isolation, and quantification

48 hours prior to transfection, AAV-293 cells (Cell Biolabs Inc) were seeded at a density of 2×105 cells/cm2. When cells reached 70-80% confluency they were transfected using a reaction mix containing pRV1 (AAV2) (37.5 µg), pH21 (AAV1) (37.5 µg), pFdelta6 (Ad Helper) (150 µg), one of the pAAV constructs described above (75 µg), pEGFP-N3 (25 µg), CaCl_2_ (150 mM) and HEPES buffered saline (HBS). The medium was replaced after 6h. 72 hours after transfection, cells were scraped in cold PBS and collected in reaction tubes. Cells were centrifuged at 800 x g for 10 minutes, cell pellets were resuspended and incubated for 1h at 37°C in a buffer containing 20 mM Tris, 159 mM NaCl (pH 8.0), 0.5% sodium deoxycholate and 50U/ml Benzonase. After two centrifugation steps at 3000 x g to remove cell debris the AAV was isolated from the supernatant using heparin columns (GE Healthcare). After loading the column with the virus particles and washing with buffers containing 20 mM Tris and increasing concentrations of NaCl (100 mM, 200 mM, 300 mM), the virus was eluted using buffers containing 20 mM Tris and 400 mM, 450 mM and 500 mM NaCl. Virus was then concentrated using Amicon Ultra-4 centrifugal filter units (Merck-Millipore) and the solution was filtered with a 0.2 µm Acrodisc syringe filter (Sigma-Aldrich).

For virus quantification, the virus load was determined using quantitative real-time PCR (qPCR). As a standard, the AAV plasmid construct was used to generate a dilution series with known concentrations. The diluted plasmid and the solutions containing AAV in an unknown concentration have been used as template for a PCR amplifying the Woodchuck Hepatitis Virus Posttranscriptional Regulatory Element (WPRE) using the oligonucleotides WPRE-F 5’-TGCTTCCCGTATGGCTTTCAT-3’ and WPRE-R 5’-CAGCAAACACAGTGCACACC-3’ as primers and SYBR Select Master Mix (ThermoFisher Scientific). The measurements were performed with a CFX384 instrument (Biorad) and analyses were done using the CFX Manager software (Biorad).

### Immunohistochemistry, dendritic spine analysis, and protein analysis

Isoflurane-anesthetized mice were transcardially perfused with PBS and subsequently with 4% PFA (for immunohistochemistry). The detailed procedures for tissue preparation, immunohistochemistry, Golgi staining and dendritic spine analysis of CA1 pyramidal neurons, Western Blot and capillary Western immunoassay are described in Supplementary material and methods.

### Analysis of Gene-expression and regulatory networks

#### Probe sampling

Isoflurane-anesthetized mice were sacrificed by decapitation and brains were harvested. Brain hemispheres were snap frozen in liquid nitrogen and brain tissue was stored at −80°C until further use.

#### RNA Isolation

Total RNA extraction was carried out using QIAzol lysis reagent and RNeasy Mini Kit (Qiagen, Hilden, Germany) according to manufacturer instructions; quality control of isolated RNA was performed on a QIAxcel capillary electrophoresis system (Qiagen, Hilden, Germany). Integrity of RNA was proven by reporting the 28S/18S ratio and the RIS number (RNA Integrity Score) for analyzed samples. Quantification of total RNA and measuring the A260/A280 ratio was performed on a Nano-Drop spectrophotometer ND-2000 (Thermo Fisher Scientific, Schwerte, Germany).

#### Microarray Analysis

250 ng of total RNA of each sample was reversely transcribed and amplified using the TargetAmp-Nano Labeling Kit for Illumina Expression BeadChips (Biozym Scientific GmbH, Hessisch Oldendorf, Germany) on a Biorad MJ Thermal Cycler (Biorad, Munich, Germany) according to the manufacturer’s instructions. All cRNA samples were purified with the NucleoSpin RNA Clean-up system (Macherey-Nagel, GmbH & Co. KG, Düren, Germany) and quantified on a Nano-Drop spectrophotometer ND-2000 (Thermo Fisher Scientific, Schwerte, Germany) before hybridization. Samples (750 ng cRNA) were hybridized 20 h, 58°C on two Illumina MouseRef-8 v2.0 Expression BeadChips (Illumina, San Diego, CA, USA). Post-hybridization data read-out, data pre-processing including spot detection, control probe quality check, background subtraction, signal averaging, and data analyses were performed with the Illumina GenomeStudio-Software v.2011.1 (Illumina, San Diego, CA, USA).

#### Data processing, gene set enrichment analysis and analysis of regulatory networks

Microarray data were preprocessed in the R statistical environment using the lumi package,^38^ using log_2_-transformation and robust spline normalization (rsn). One sample (“9298740075_G”) was identified as an outlier based on hierarchical clustering and was removed (Supplementary Fig. 2, A and B). Differentially expressed genes between PCI and Sham were determined using FDR/Benjamini-Hochberg correction and log_2_FC ≥ 1 cut-off. Gene set enrichment analysis for enriched Gene Ontology (GO) terms (category biological pathway) was performed using the piano package.^39^ The raw and pre-processed data have been deposited in NCBI GEO DataSets with the following accession number: GSE167610.

Microarray data were further processed using the R environment (R Studio Version 1.1.383, R Version 3.6.1; RStudio Team, RStudio, Inc., Boston, MA, R Core Team, Foundation for Statistical Computing, Vienna, Austria) with the packages reshape2^40^ and ggplot2. Networks were built and analyzed with the aid of Cytoscape (Version 3.8.0) including the apps Legend Creator (version 1.1.5) and stringApp (version 1.6.0).^41^ Protein interactions were imported by the stringApp from the string data base for the species Mus musculus with a confidence cut-off ≥0.4.

### Statistical analysis

Our presented data represent the median or the mean ± SEM as indicated in the figure legends. Depending on normal distribution and variance of data an unpaired, 2-sided Student’s t test or a 2-way ANOVA (> 2 groups) or non-parametric tests were used to evaluate statistical significance using P less than 0.05 as the cutoff.

### Data availability

The raw and preprocessed Microarray data have been deposited in NCBI GEO DataSets with the accession number GSE167610. All further data are available in the main text or the Supplementary Materials.

## Results

### Experimental sepsis causes neurocognitive dysfunction, defective synaptic plasticity, and synaptic scaling in excitatory synapses

Behavioral Observations: We induced severe experimental sepsis in a total of 769 male C57BL/6J mice by polymicrobial peritonitis after intraperitoneal injection of a standardized and microbiologically validated human faeces material (peritoneal contamination and infection model, PCI; Fig. 1A)^27^. Mice showed a median disease severity of 12.3 on the cumulative 5 day PCI Clinical Severity Score (CSS; Fig. 1B) and only surviving mice with a 5 day CSS of ≥ 11.5 indicating severe sepsis were included for further experiments (n = 309 PCI). Concomitant antibiotic rescue therapy with meropenem (weeks 1 and 2) and enrofloxacin from week 3 on led to a survival rate of 0.38 after 8 weeks (Fig. 1B). At a late stage, 8 weeks after PCI induction, the physically recovered mice still showed delayed learning and defective memory performance as revealed by Morris water maze test (MWM). Deficits were present over the complete learning phase in the MWM and memory dysfunction persisted also into the recall phase on day 8 (Fig. 1C). Moreover, mice had increased anxiety-like behavior in the elevated plus maze test (EPM, Fig. 1D) and in the open field (OF, Fig. 1E). Since the overall phenotype was already normalized as indicated by recovered body weight and regular locomotor activity, the behavioral abnormalities of defective memory and increased anxiety can be attributed to inflammation-induced brain dysfunction and not to post-sepsis related general sickness (Fig. 1, D and E).

**Figure. 1.**
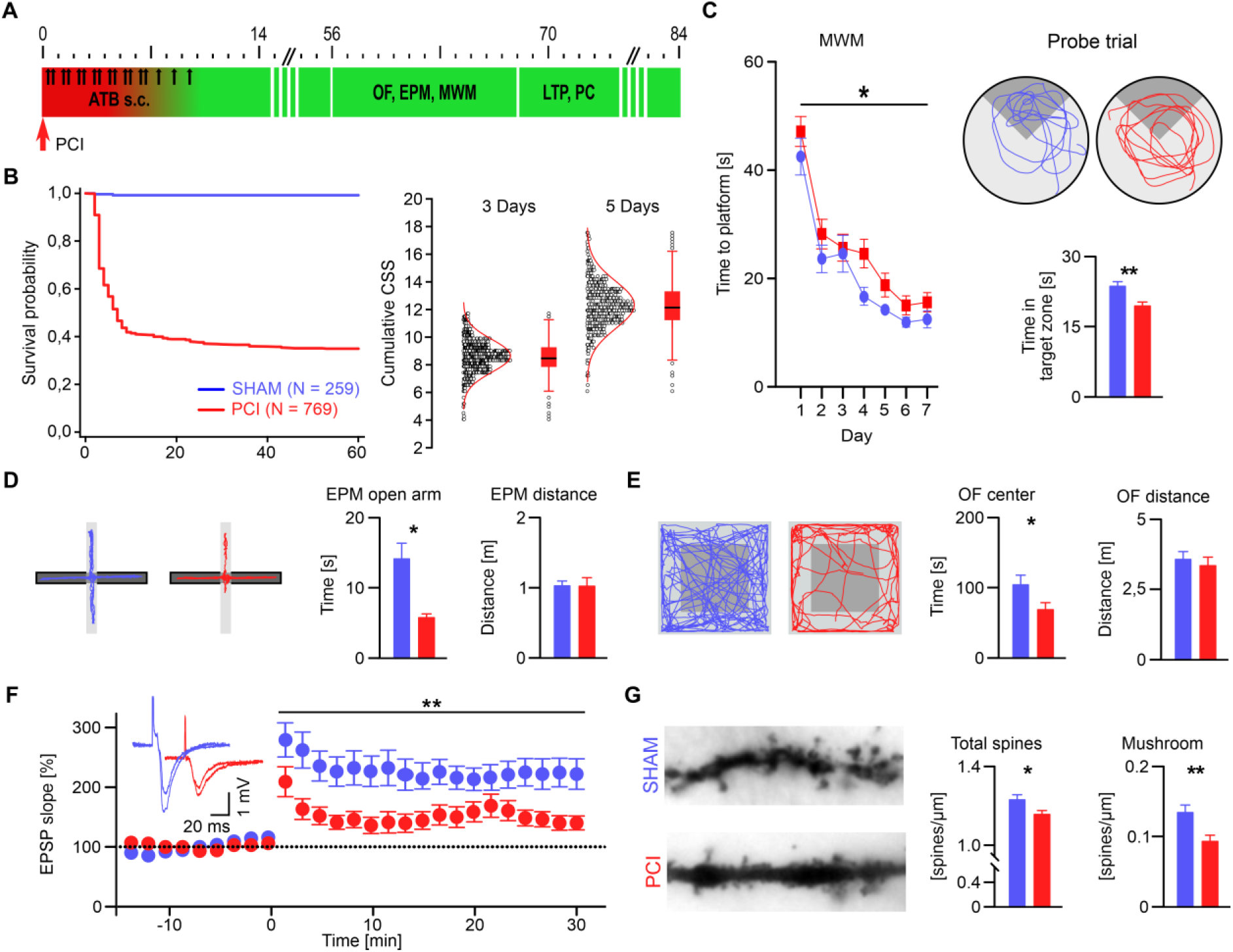
Experimental sepsis causes neurocognitive dysfunction, defective long-term potentiation and reduced neuronal spine density. (**A**) Experimental schematic: acute phase of PCI is marked in red. Antibiotics (ATB) are delivered i.p. until day 10 and behavioral testing (open field, OF; elevated plus maze, EPM; Morris water maze, WMW) and *ex-vivo* electrophysiological recordings (long-term potentiation, LTP, and patch-clamp recordings, PC) are performed at the indicated time points. (**B**) Kaplan-Meier survival curve of experimental animals (left). Mice intentionally leaving the study e.g. for tissue harvesting were included up to the time when they are censored. Mean and distribution of sepsis severity score (CSS) on day 3 and on day 5 after PCI (right; each circle represents an individual mouse after PCI). (**C**) In the MWM PCI mice (n = 12, red) show affected learning (left; 2-way repeated measures ANOVA) and reduced memory recall in the probe trial (right; example tracking traces are shown, target quadrant in dark grey; SHAM mice n = 18, blue). (**D**) PCI mice spent reduced time in the EPM open arms (light grey). The locomotor activity as measured by total distance traveled within the EPM is unchanged (n = 12/7; SHAM/PCI). (**E**) PCI mice spent reduced time the OF center (dark grey), whereas the distance traveled within the OF as a measure for locomotor activity is unchanged (n = 20/15; SHAM/PCI). (**F**) Long-term potentiation (LTP) in the hippocampal Schaffer collateral – CA1 pathway is severely impaired in PCI mice (n = 7; SHAM: n = 12; 2-way repeated measures ANOVA). Inset shows average traces of example recordings before and after theta burst stimulation. (**G**) Density of total synaptic spines and of mature mushroom spines in apical dendrites of CA1neurons is reduced after PCI (n = 42/33). Bar graphs show mean ± SEM; 2-tailed Student’s t test, unless otherwise indicated. *P < 0.05; **P < 0.01.

Synaptic function and plasticity: In order to identify the underlying pathophysiology of cognitive dysfunction we tested synaptic plasticity by recording local field potentials in the CA1 region of the hippocampus after stimulation of Schaffer collaterals. We found a severely reduced long-term potentiation (LTP) at week 10 after PCI induction (Fig. 1F). To substantiate these functional defects, we additionally quantified synaptic spines in apical dendrites of hippocampal CA1 neurons providing morphological information on plasticity. The overall spine density and, more specifically, mature mushroom spines were reduced corroborating the findings of defective LTP (Fig. 1G). Together, these data indicate disturbed synaptic plasticity as the potential basis for persistent cognitive decline in the late phase after surviving sepsis.

Next, we investigated synaptic transmission in the hippocampus from week 10 after PCI by whole-cell patch-clamp recordings in dentate gyrus granule cells. We found a reduced frequency of quantal excitatory postsynaptic currents (miniature, mEPSCs, Fig. 2A) and spontaneous EPSCs (sEPSCs, Fig. 2B) which may indicate overall reduction of excitatory presynaptic terminals. Unexpectedly, peak amplitudes of mEPSCs, sEPSCs, and of minimally evoked EPSCs (i.e. eEPSCs arising from individual synapses) were increased after stimulation of the medial perforant pathway (Fig. 2A to C) whereas EPSC rise and decay time kinetics were unchanged (Supplementary Fig. 1). This might reflect increased recruitment of AMPA receptors to postsynaptic sites suggesting another mode of inducing gain in plasticity in a homeostatic scaling mechanism.

**Figure 2.**
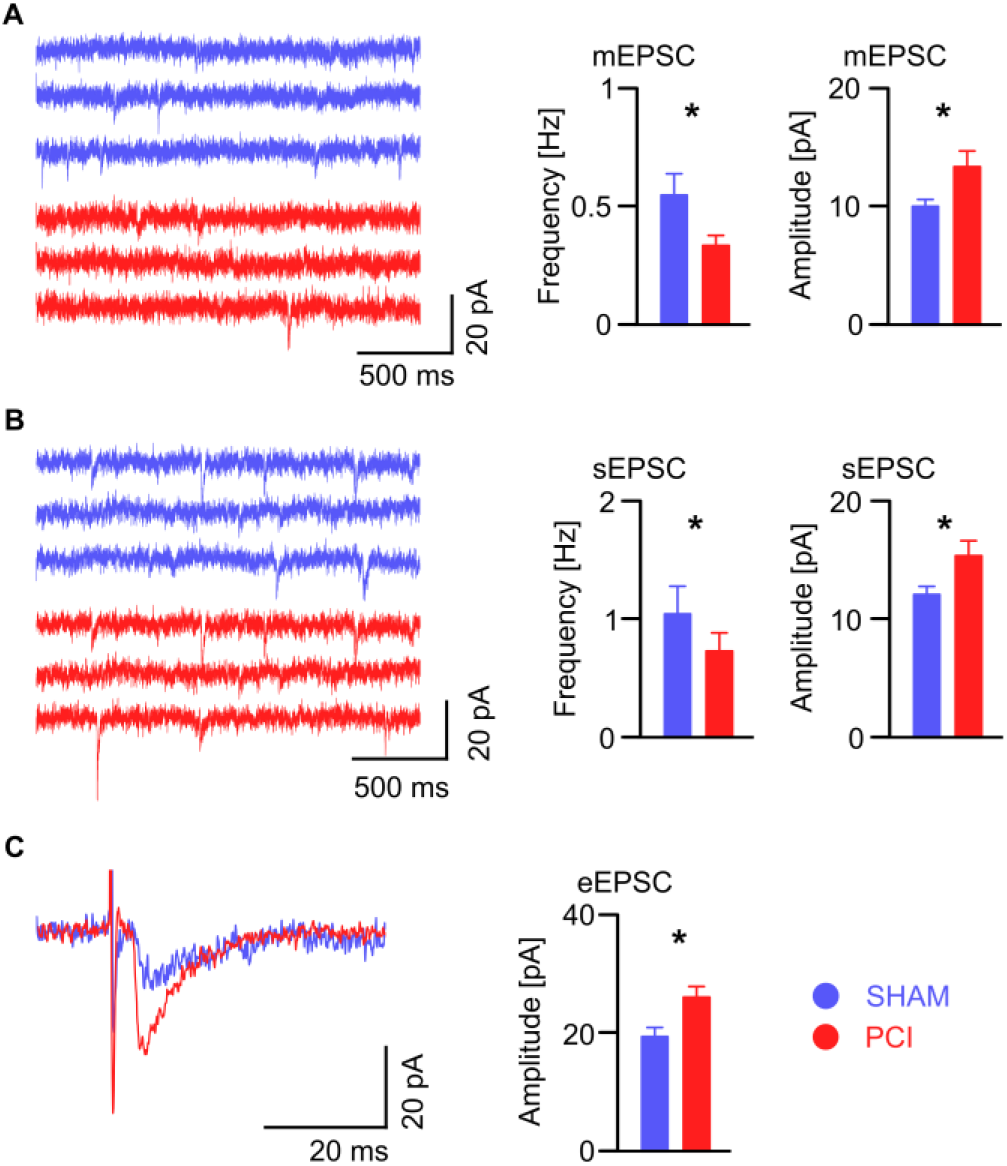
Synaptic scaling of excitatory synaptic transmission in mice after sepsis. Patch-clamp recordings of dentate gyrus granule cells indicates reduced frequency of (**A**) miniature (m) and (**B**) spontaneous (s)EPSCs, and (**C**) increased peak amplitude of minimally evoked (eEPSCs). mEPSCs, sEPSCs and eEPSC (PCI: n = 11/13/13, SHAM: n = 11/13/16, respectively). Example traces in (**A**) and (**B**), average traces of minimal evoked eEPSCs in (**C**). Bar graphs show mean ± SEM; 2-tailed Student’s t test, unless otherwise indicated. *P < 0.05; **P < 0.01.

### PCI leads to distinct and time-dependent changes in neuronal pathways relevant for synaptic plasticity

In order to obtain mechanistic insights into the molecular mechanism of sepsis-induced neuronal dysfunction we next investigated the transcriptional profile of brain tissue of acute septic PCI mice (day 3 after PCI) and of mice at a very late stage after surviving sepsis (week 10), and compared this with matched SHAM control mice. We found differently regulated brain gene expression early after sepsis, whereas after 10 weeks gene expression on the level of single genes was similar to SHAM controls (Fig. 3 and Supplementary Fig. 2). Gene Ontology (GO) term enrichment analysis of differentially expressed genes (DEGs) at day 3 revealed that transcripts related to neuronal function (e.g., GO terms ‘synapse’, ‘postsynaptic density’, ‘dendrite’) were markedly downregulated (Fig. 4). As expected, transcripts related to inflammation (e.g. GO terms ‘innate immune response’, ‘inflammatory response’) and to gene transcription were all highly upregulated. In contrast, in the late stage after sepsis, inflammatory pathways in the brain were no longer activated and pathways related to gene transcription were downregulated. Moreover, genes associated to neuronal function (e.g. GO terms ‘neuron projection’, ‘dendrite’) were upregulated, indicating compensatory or maladaptive regulation once severe systemic inflammation had subsided. The most prominently downregulated gene was *Arc*, encoding activity-regulated cytoskeleton-associated protein ARC/ARG3.1 (ARC), a master regulator of synaptic plasticity ^42^ (Fig. 5, A and B). The protein interaction network of corresponding differently expressed genes on day 3 after PCI revealed downregulation of genes encoding close ARC interaction partners, thus predicting a major role in dysfunctional synaptic plasticity and signaling (Fig. 5A). Genes of the (1) early growth response protein (*Egr*) family, i.e. transcription regulatory factors involved in neuronal plasticity ^43^, (2) *Dusp6* encoding the ERK-directed dual-specificity phosphatase 6, (3) *Nrgn* encoding Neurogranin/RC3, a calmodulin-binding protein involved in CaMKII dependent synaptic plasticity ^44^, and (4) *Mapk11*, encoding p38 Mitogen-activated protein kinase 11, were all downregulated (Fig. 5, A and B). At the protein level and supporting our interpretation after PCI, we indeed found a reduction of ARC and of phosphorylated ERK, an upstream mitogen-activated protein (MAP) kinase relevant for transcription of immediate-early genes such as *Arc* ^45^ at the early stage but also 10 weeks after PCI (Fig. 5, C and D; Supplementary Fig. 3). Together, these data indicate that severe neuroinflammation goes along with alterations of synaptic regulatory pathways in early stages whereas maladaptive regulation of neuronal pathways appears in late stages. Impaired neuronal functionality as shown by electrophysiology converges on activity-regulated pathways that are relevant for synaptic plasticity.

**Figure 3.**
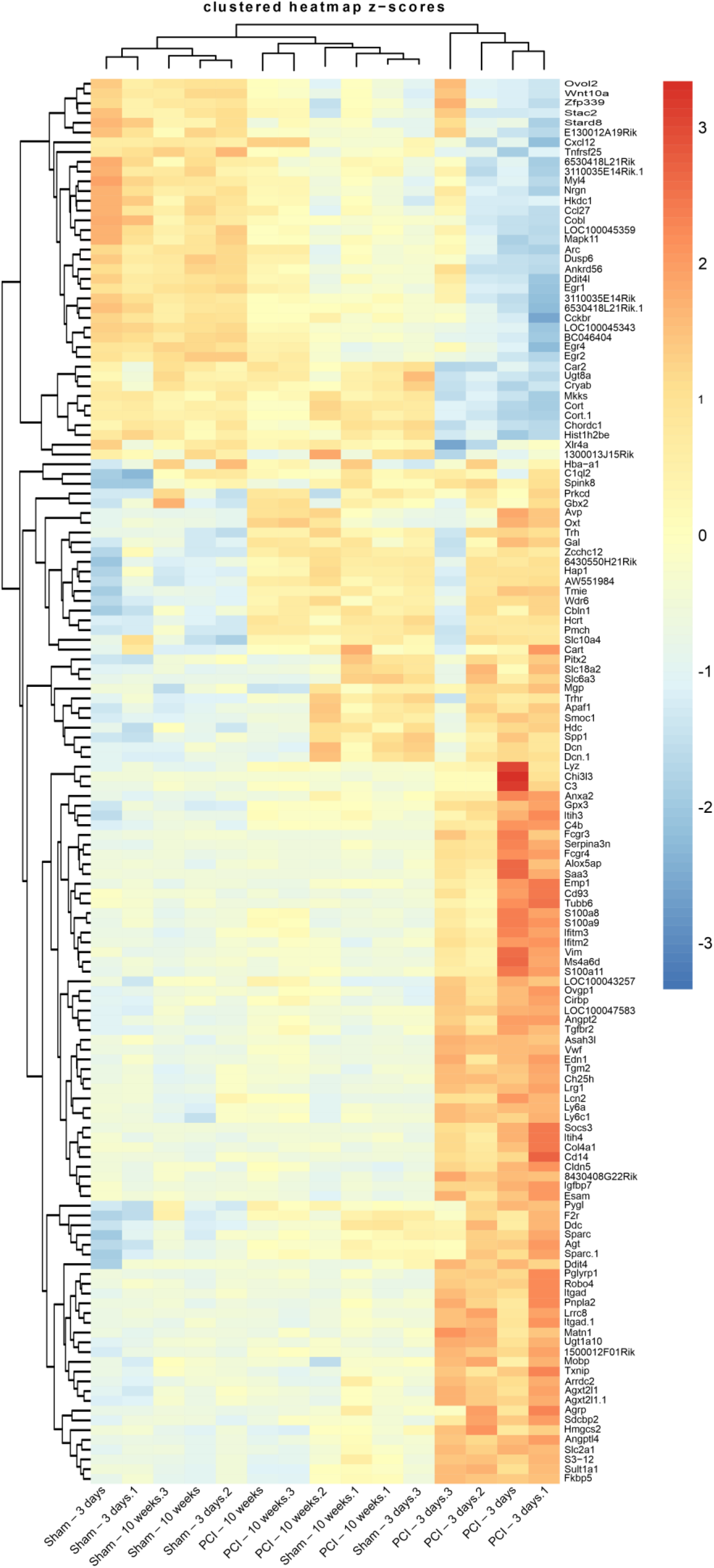
Hierarchically clustered heatmap of DEGs in a murine sepsis model. Expression of 144 DEGs (log_2_FC > 1) in brain tissue on 3 days and 10 weeks after PCI with (color-coded) normalized z-score visualization. Biological replicates from 3 days after PCI distinctly cluster together (four lanes on the right).

**Figure 4.**
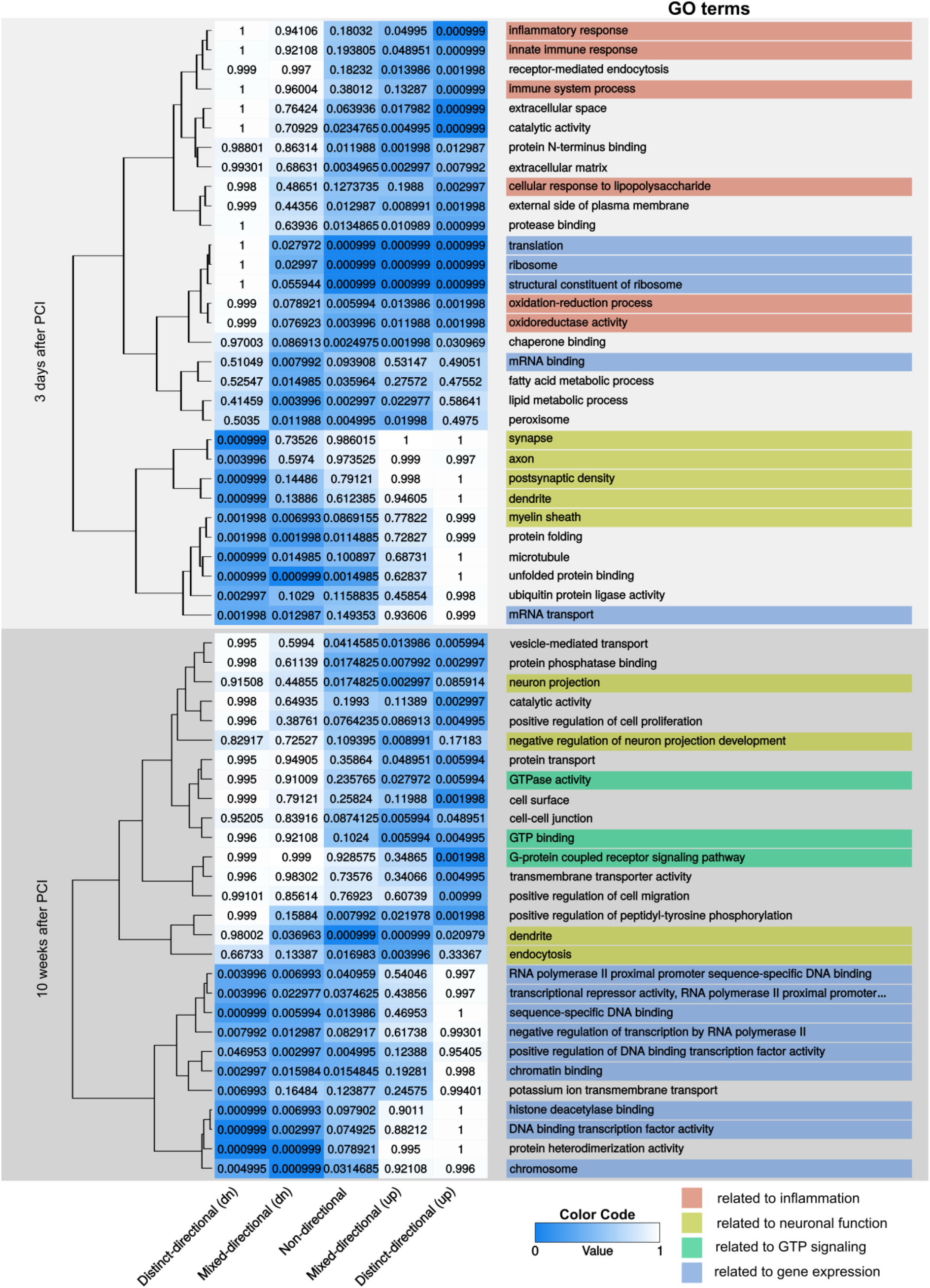
PCI leads to distinct and time-dependent changes in gene expression in the brain. Gene Ontology (GO) terms enriched in differently expressed genes (144 DEGs; log_2_FC > 1) found by microarray analysis of brain tissue at 3 days and 10 weeks after PCI. Differentially expressed genes were interrogated for enrichment in the gene sets of the GO definitions employing the piano package. Significantly enriched gene sets were clustered according to the score values of the five piano criteria distinct directional (up and down), mixed directional (up and down) and non-directional.

**Figure 5.**
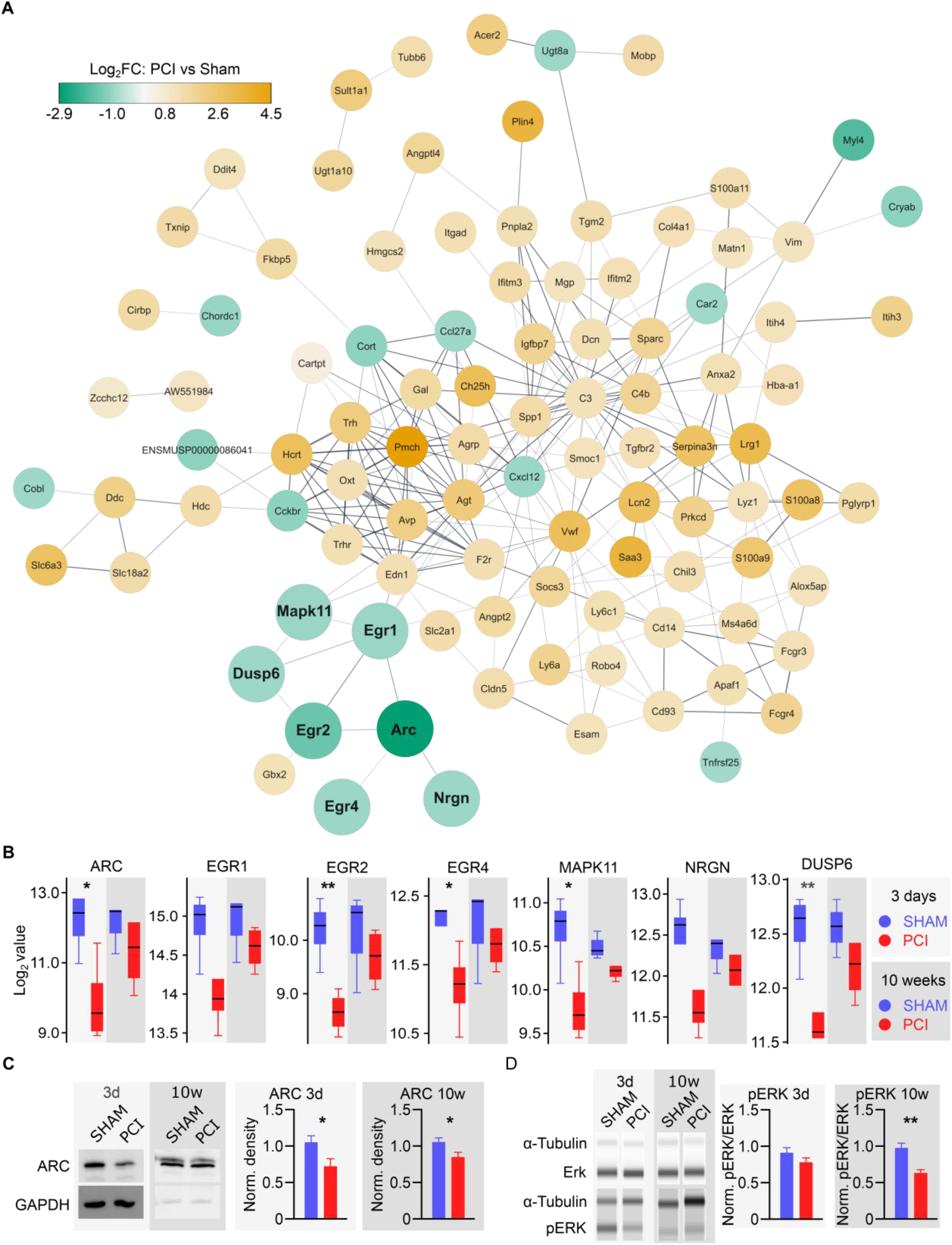
PCI induces downregulation of neuronal pathways relevant for synaptic plasticity. (**A**) Interaction network of proteins corresponding to DEGs on day 3 after PCI. The node color depicts the respective Log_2_FC from microarray data. Proteins related to ARC synaptic signalling are highlighted by size. (**B**) Expression of seven selected DEGs that are related to ARC signalling and synaptic function (highlighted in **A**). 3 days SHAM and PCI and 10 weeks PCI n = 4, 10 weeks SHAM n = 3. Boxplots show median and 75^th^ to 25^th^ interquartile range, whiskers show respective maximum or minimum values within 1.5 times of the interquartile range; 2-tailed Student’s t test. (**C** and **D**) Quantitative analysis of hippocampal ARC protein and pERK/ERK protein ratio (n = 8, each; normalized to SHAM ARC or SHAM pERK/ERK ratio, respectively) shows downregulation of signalling pathways in the network shown in (**A**). 2-tailed Student’s t test. *P < 0.05, **P < 0.01.

### Rescue strategies of defective plasticity

Based on these findings, we then followed two interventional strategies directed at rescuing defective synaptic plasticity and cognitive dysfunction after PCI. First, we specifically targeted the PCI-induced downregulation of the synaptic master regulator ARC by overexpressing *Arc* using a neuron-specific adeno-associated virus. Secondly, in a completely different and systemic behavior-modulating paradigm we subjected mice to enriched environment (EE) which is known to have beneficial effects for cognitive performance by enhancing synaptic plasticity. The underlying mechanisms for these effects are thought to be initiated by upregulation of activity-regulated genes and pathways ^46,47^.

### Overexpression of *Arc* rescues sepsis-induced cognitive deficits and impaired long-term potentiation

We found increased AMPA receptor mediated EPSC peak amplitude in synaptic transmission in mice after PCI (Fig. 2, A to C) which is consistent with the observation that deficiency of ARC leads to upscaling of AMPA receptor expression ^48^. To overcome the identified basal and not only activity-dependent PCI-induced synaptic downregulation of ARC we constitutively overexpressed *Arc* in the hippocampus of post-septic mice: we here used an adeno-associated virus containing a neuron-specific bicistronic plasmid of the *Arc* and *Venus* transgene under the synapsin promotor (*Arc*-AAV; Fig. 6A). Bilateral *in-vivo* injection of *Arc*-AAV in three injection sites of the hippocampus led to a robust, constitutively driven and not activity-dependent upregulation of ARC in hippocampal neurons (Fig. 6B). According to previous studies showing inverse regulation of synaptic AMPA receptor expression by synaptic scaling mechanisms after *Arc* overexpression ^49–51^ we expected a diminished LTP due to disturbed AMPA receptor mediated signaling in SHAM mice after *Arc* overexpression. Indeed, we found reduced LTP in the Schaffer collateral-CA1 pathway in patch-clamp single cell recordings of SHAM mice after *Arc*-AAV injection (Supplementary Fig. 4).

**Figure 6.**
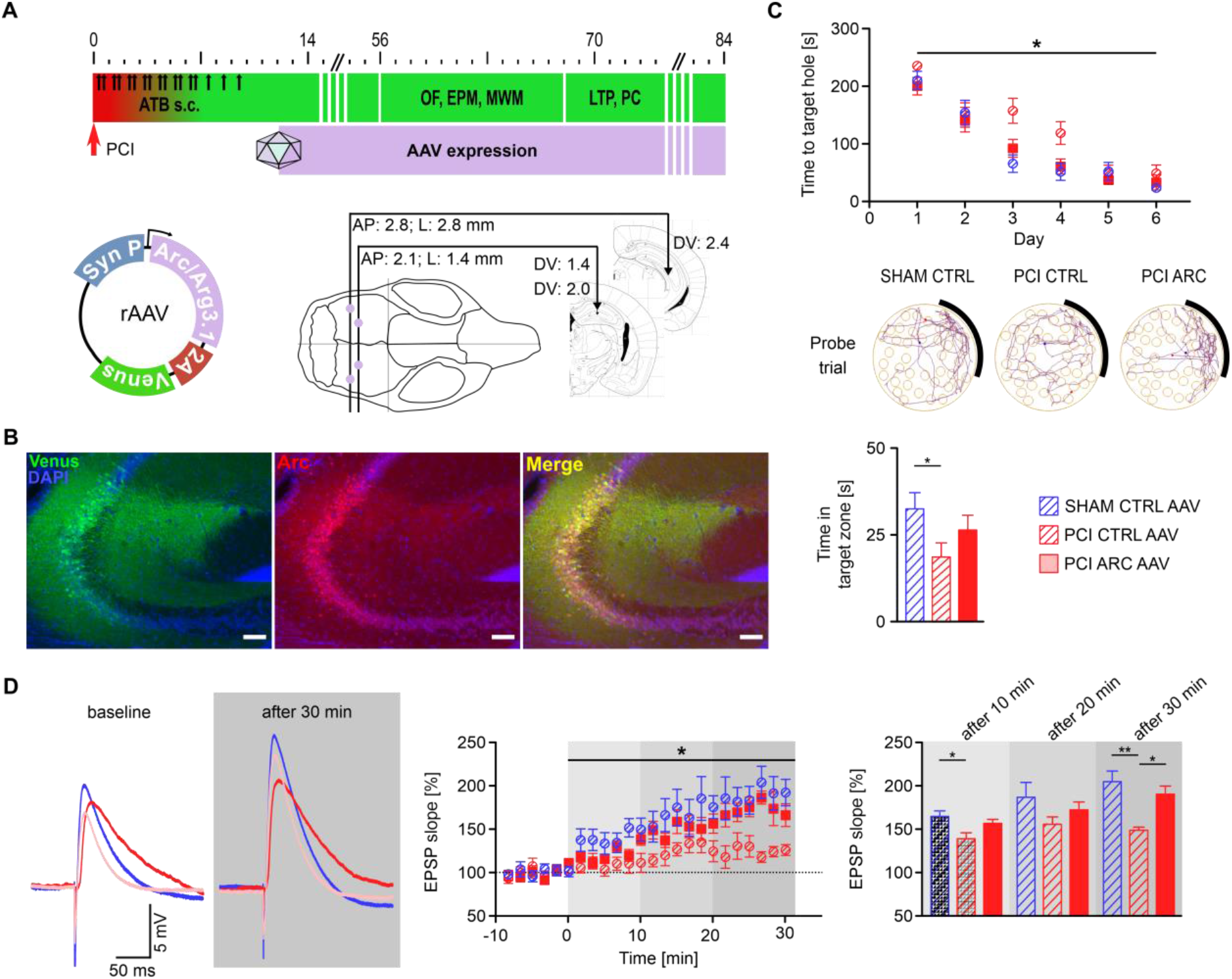
Hippocampal overexpression of *Arc* rescues inflammation-induced cognitive dysfunction and defective long-term potentiation. (**A**) Experimental schedule indicating ARC overexpression in the hippocampus from day 10 after PCI. Stereotactic intrahippocampal microinjection of an adeno-associated virus (AAV) containing the bicistronic vector for ARC overexpression and Venus fluorescence under the synapsin promotor at the indicated coordinates. (**B**) Representative example of constitutive ARC overexpression in the CA3 region of the hippocampus. Scale bars: 40 μm. (**C**) PCI mice receiving control AAV (PCI CTRL AAV; n = 17) show learning deficits in the Barnes Maze compared to SHAM CTRL AAV (n = 18) as indicated by prolonged time to the target hole (upper panel; 2-way repeated measures ANOVA) and reduced time in the target zone in the probe trail (day 7, lower panel). ARC overexpression in PCI mice (PCI ARC AAV; n = 18) results in rescue of learning and memory dysfunction. Example traces indicate tracks of individual mice during the probe trial; the target quadrant containing the open hole is marked in grey. (**D**) LTP as measured by slope of excitatory postsynaptic potentials (EPSPs) in single-cell whole cell recordings in CA1 neurons after stimulation of Schaffer collaterals. Paired stimulation results in LTP in SHAM CTRL AAV (n = 11) and PCI ARC AAV (n = 8) mice whereas PCI mice with control virus injection (PCI CTRL AAV; n = 9) show reduced LTP. Left panel: average traces (blue: SHAM CTRL AAV; red: PCI CTRL AAV; light red: PCI ARC AAV). Middle panel: LTP time course. Right panel: comparison of LTP at the indicated time points after theta burst stimulation.; 2-way repeated measures ANOVA. *P < 0.05, **P < 0.01.

PCI surviving mice that received injection of the empty control vector expressing only fluorescent Venus protein again showed learning and memory dysfunction also in the Barnes maze (BM) 8 weeks after PCI, similar to the PCI induced alterations in the Morris water maze (MWM, Fig. 6C). In contrast, hippocampal *Arc* overexpression in PCI mice induced improvement of spatial learning and memory recall (Fig. 6C). As expected, *Arc* overexpression in SHAM mice did not improve cognitive function (Supplementary Fig. 5). Locomotor function was not affected in all experimental groups (Supplementary Fig. 6).

Fear-related behavior as measured in the EPM was present in PCI mice and did not improve upon *Arc* overexpression in the hippocampus since it is primarily mediated by signaling pathways in the amygdala (Supplementary Fig. 6B). Corroborating the behavioral findings, hippocampal LTP of PCI mice was rescued after *Arc*-AAV injection to almost normal values when compared to SHAM mice having received injections of *control*-AAV (Fig. 6D). These data suggest that increasing ARC protein levels in the hippocampus may be effective to in part reverse sepsis-induced neuronal dysfunction and defective synaptic plasticity mechanisms, relevant for cognitive functions.

### Enriched environment improves synaptic pathology and cognitive dysfunction induced by PCI

Once we had established that activity-induced synaptic signaling genes and proteins are severely downregulated after PCI, we used a second, completely different paradigm to address the hypothesis that cognitive dysfunction in post-sepsis mice may also profit from stimulus-induced general activation. We therefore utilized an established EE paradigm ^46^ to investigate its rescuing potential for defective synaptic plasticity and memory dysfunction induced by PCI (Fig. 7A). As a first step, we again reproduced PCI-induced long-term cognitive dysfunction, anxiety-related behavior, and defective LTP in a separate cohort of mice without special housing conditions (PCI versus SHAM non-enriched (NE); Fig. 7, B to D). Indeed, when PCI mice were subjected to 6 weeks of EE housing conditions, they showed improved performance in the BM test 8 weeks after PCI and reduced anxiety-related behavior in the EPM (Fig. 7, B and C). Possible confounders such as PCI-induced sickness with locomotor hypoactivity were no more present in any of the surviving mice in the experimental groups at 8 weeks after PCI (Supplementary Fig. 7). Synaptic plasticity as measured by hippocampal LTP was equally preserved and reduction of synaptic mushroom spines was no longer present in PCI mice following this EE paradigm (Fig. 7, D and E).

**Figure 7.**
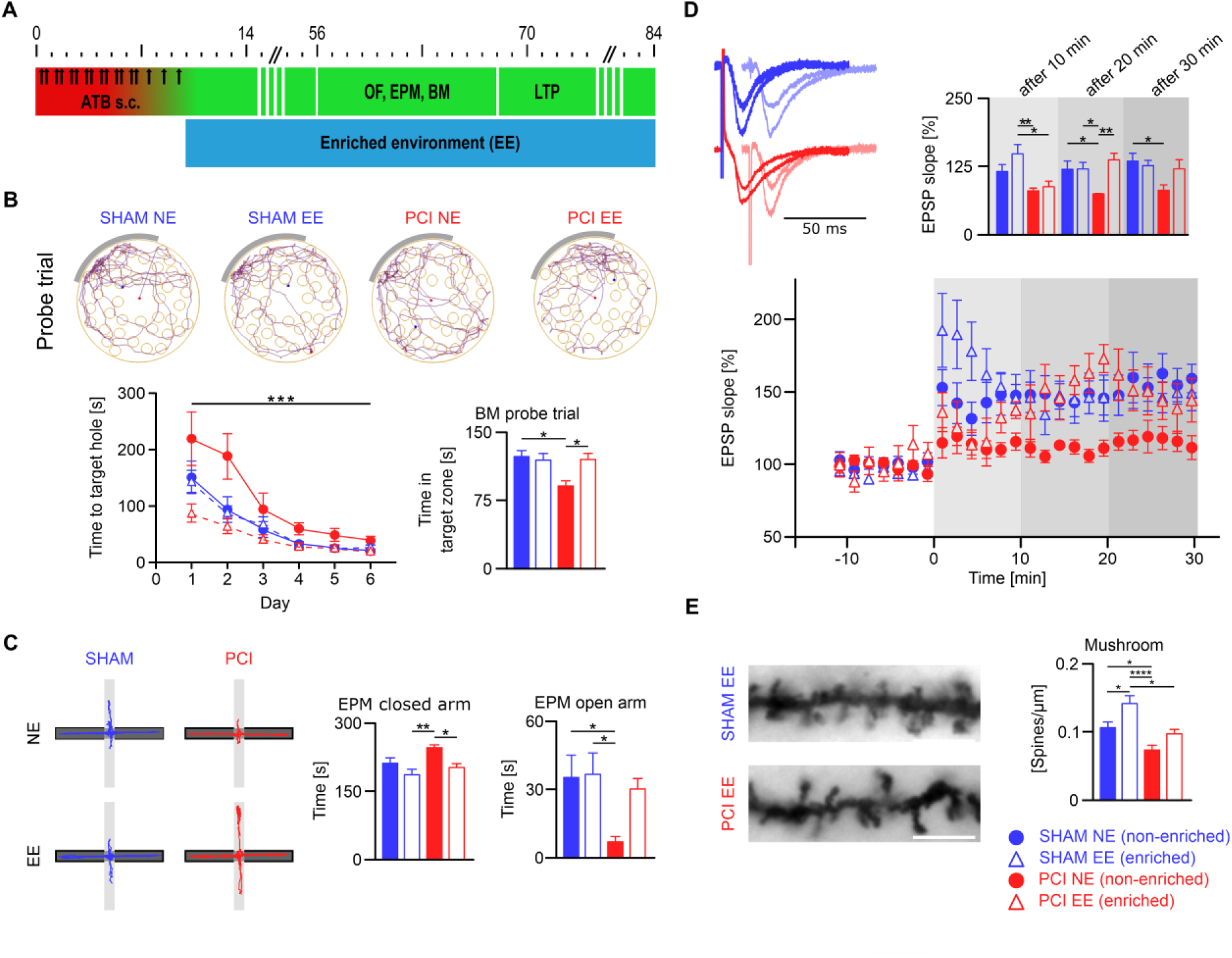
Enriched environment leads to improvement of synaptic pathology and cognitive dysfunction after PCI. (**A**) Mice were subjected to enriched environment (EE) or standard housing (NE) from day 10 after PCI. (**B**) EE improves cognitive dysfunction in the BM after PCI (PCI EE) as indicated by reduced time to target hole in the training phase (left;; 2-way repeated measures ANOVA) and by increased time in the BM target zone during the probe trial compared to PCI mice in standard care (PCI NE). Example traces indicate tracks of individual mice during the probe trial; target quadrant is marked in grey. (**C**) EE leads to reduced anxiety-related behavior in the EPM after PCI as shown by reduced time in the closed and increased time in the open arms (n = 14 / 18 / 13 / 16; SHAM NE / SHAM EE / PCI NE / PCI EE in B and C). (**D**) LTP as evaluated by field potential recording in CA1 is improved in PCI mice after EE. Left panel: average traces before and after theta burst stimulation (SHAM EE light blue; PCI EE light red; amplitudes of the baseline traces of each group were scaled to the amplitude of the SHAM NE trace (dark blue). Right panel: comparison of LTP at the indicated time points after theta burst stimulation; bottom panel: LTP time course. (E) Quantification of synaptic mushroom spines in apical dendrites of CA1 neurons. Example images are provided (scale bar: 5 µm). PCI leads to reduction of mature mushroom spine density (PCI NE) which is rescued to basal levels (SHAM NE) after enriched environment (PCI EE); n = 42 / 21 / 33 / 36; SHAM NE / SHAM EE / PCI NE / PCI EE.; 2-way repeated measures ANOVA. *P < 0.05, **P < 0.01, ***P < 0.001.

## Discussion

From our findings we can define three major aspects that may help to understand synaptic pathology after severe inflammation. Any of these may serve as targets for rescue strategies of disordered synaptic function and eventually help to improve inflammation-induced cognitive decline. First, we identified defective synaptic plasticity as a crucial pathogenic mechanism of long-term cognitive dysfunction in SAE persisting beyond acute brain inflammation ^52^. More specifically, we uncovered distinct changes in the regulation of synaptic proteins leading to dysfunctional basal synaptic transmission and scaling and defective LTP. These synaptic changes developed in early stages of SAE in parallel to severe sepsis-induced neuroinflammation and persisted even when the acute inflammation had subsided. Second, the induced overexpression of severely downregulated ARC, a prominent mediator of synaptic signaling, appears as a crucial and sufficiently effective mechanism to largely rescue sepsis-induced synaptic and brain dysfunction. Third, SAE-induced synaptic changes and memory dysfunction could also be restored by non-invasive activation measures such as EE.

Several studies reported the effects of severe systemic inflammation on immune mechanism in the brain. Systemic application of lipopolysaccharide (LPS) results in early inflammation-associated alterations and activation of cerebral endiothelial cells ^10^. This initial event is followed by glial cell activation, e.g. of microglia, thus leading to an inflammatory state in the CNS characterized by increased production and release of proinflammatory cytokines and chemokines ^15,53^ including tumor necrosis factor-α (TNFα), interleukin-1β (IL-1β), IL-6, and CXCL10. In parallel, anti-inflammatory mediators, e.g. lipocalin2 (LCN2) are induced to counteract LPS-induced neuroinflammation ^16,54,55^. Even if experimental systemic LPS challenge may not completely mimick polymicrobial sepsisthese observations likely reflect important mechanistic aspects of CNS neuroinflammation since this proinflammatory profile has also been identified in other infectious rodent models, e.g. in the cecal ligation and puncture (CLP) model ^56^.

Direct interactions of proinflammatory mediators with synaptic function and excitatory signaling are well established. Endogenous TNFα has been shown to enhance the membrane insertion of synaptic AMPA receptors leading to increased EPSC amplitude ^57^ and to synaptic scaling. Moreover, proinflammatory cytokines are able to inhibit LTP via disturbed mitogen-activated protein kinase (MAP-K) signaling ^58,59^ and increased cytokine levels are associated with cognitive decline ^60^. Here, we identified dysregulation of ARC together with its close interaction partners and the MAPK downstream mediator ERK which probably underlies dysregulation of excitatory synaptic signaling. *Arc* is an immediate early gene (IEG) and its transcription and translation are directly regulated by neuronal activity ^61^. ARC is abundantly expressed in cell bodies and dendrites of CamKII positive excitatory neurons in the CNS ^62^. Although recent studies suggest that ARC is not indispensable for LTP ^63^, it is thought to be critical for memory formation and an important, bidirectional regulator of long-term synaptic plasticity including LTP, long-term depression (LTD), heterosynaptic LTD, and homeostatic synaptic scaling ^42^. ARC mediates activity-regulated trapping of AMPA receptors at postsynaptic sites by interaction with the endocytic machinery and PSD-95 complexes ^64^. Indeed, in *Arc* knockout (ko) mice and after *Arc* silencing using antisense oligonucleotides memory formation and consolidation of LTP is impaired ^65,66^. Furthermore, ARC regulates synaptic AMPA receptor expression dependent on synaptic activity, representing the key mechanism of homeostatic plasticity ^67,68^. In our PCI model, we identified substantially reduced *Arc* transcripts and protein levels already under basal conditions without additional external stimulation. Accordingly, as one might expect from *Arc* ko studies, we found increased AMPA receptor currents in basal conditions and LTP was severely affected. Our experiments in mice with constitutive overexpression of *Arc* therefore aimed at rescuing the sepsis-induced basal and persistent downregulation of ARC. This strategy was chosen following the observations by others that *Arc* overexpression in cell culture and in organotypic brain slices results in blockade of homeostatic upregulation of AMPA receptors and reduction of EPSC amplitude ^48,50^. Our present findings with overexpression of *Arc in-vivo* show that this approach was successful in rescuing PCI-induced hippocampal dysfunction thus providing evidence for a successful strategic intervention in a crucial pathway of disturbed synaptic signaling.

Next, we questioned whether a completely different approach of general activation which also upregulates activity-dependent pathways might induce beneficial effects on synaptic plasticity and memory in SAE. Utilizing an established EE paradigm we were able to counteract inflammation-induced defects of synaptic plasticity and memory dysfunction. EE has been proven to be effective in various models of neurodegenerative disorders, e.g. for Alzheimeŕs disease, Huntingtońs disease and others (for review see ^47^). In these models, EE results in enhanced learning and memory and delayed disease progression ^69^. Among a variety of mechanisms, EE can enhance adult neurogenesis and network incorporation of newborn neurons ^70^, increase the expression of signaling molecules, induce synaptic plasticity pathways ^71^ and increase the number of dendritic spines, finally leading to more efficient use of neuronal networks ^72^. Mechanistically, EE enhances neurotrophic signaling, in particular of brain-derived neurotrophic factor (BDNF) via the trkB receptor and downstream activation of MAPK, e.g. the ERK pathway ^73^. Moreover, EE increases synaptogenesis and fosters synaptic plasticity like LTP ^74^, which collectively may explain the beneficial effects of EE on cognitive function observed in our present study. As shown in other models, EE can directly and extensively upregulate *Arc* ^75^ and other IEGs, e.g. *Egr 1, 2, 4* und *Dusp 6* in the hippocampus ^69,76,77^. Taken together, EE obviously acts via multiple synergistic mechanisms, and the effects on IEGs and the MAP-K pathway are plausible candidates to explain the beneficial impact on synaptic function, plasticity, and on memory after severe systemic inflammation.

Previous studies reported a time-dependent improvement of cognitive dysfunction in rats after CLP, another rodent model reflecting polymicrobial sepsis ^78^. In our study, we showed memory dysfunction in several independent experimental groups and with different standardized behavioral readouts. We cannot rule out that cognitive dysfunction is even more pronounced in earlier stages after PCI but this might always be superimposed by the general sickness induced by PCI. For this reason, we performed behavioral testing in surviving mice at a late time point (8 weeks) when systemic signs of sepsis had largely subsided.

From a translational perspective, SAE-induced cognitive dysfunction becomes increasingly relevant. Due to improved health care and intensive care medicine more often patients survive severe systemic infections and sepsis. The majority of these patients are afflicted with SAE. Studies involving large patient cohorts provided robust evidence for the increased risk of developing dementia after severe systemic inflammation ^1^. Post-infectious long-term cognitive dysfunction results not only in major disability of afflicted patients but causes also an immense burden on primary caregivers and the health care system ^2^. The high clinical impact and socio-economic importance warrants a better understanding of SAE pathophysiology for development of target-directed therapeutic strategies ^79^.

We suggest that the murine model we investigated here may reflect residual and chronic brain dysfunction in patients surviving severe inflammation. Our findings may help to elucidate SAE-induced neuronal pathomechanisms at the molecular and cellular level. Targeting synaptic pathways offers conceptual advances for translational treatment strategies in SAE. In practical terms, a general activation scheme equivalent to the EE paradigm, e.g. by personalized cognitive training approaches early in the course of SAE, may become the first to be studied in prospective randomized and controlled clinical trials. Our findings may also encourage future research addressing regulatory pathways in patients with SAE by means of targeted intervention strategies. From animal models like the ones reported here various strategies may become feasible such as increasing ARC levels, e.g. by stimulation of upstream pathways such as neurotrophin BDNF receptor/TrkB activation, ERK signaling, or CREB phosphorylation ^49,80–82^. Either of these may be promising starting points for target-directed therapies in future pilot trials.

## Abbreviations

AAV: adeno-associated viruses;
BDNF: brain-derived neurotrophic factor
aCSF: artificial cerebrospinal fluid
Arc/Arg3.1: activity-regulated cytoskeleton-associated protein
AP: anterior-posterior
BBB: blood brain barrier
BM: Barnes maze
CLP: cecal ligation and puncture
CREB: cAMP response element-binding protein
CSS: Clinical Severity Score
CXC: C-X-C motif chemokine ligand
DEG: differentially expressed genes
Dusp6: dual-specificity phosphatase 6
DV: dorso-ventral
L: lateral
EE: enriched environment
EGR: early growth response protein
EPSC: excitatory postsynaptic currents
EPM: elevated plus maze test
ERK: extracellular-signal regulated kinase
fEPSP: field Evoked Postsynaptic Potential
GO: Gene Ontology
HBS: HEPES buffered saline
IEG: immediate early gene (IEG)
Il: interleukin
LPS: lipopolysaccharide
LCN2: lipocalin 2
LTP: long-term potentiation
NE: not enriched
Nrgn: Neurogranin
Mapk11: mitogen-activated protein kinase 11
MWM: morris water maze test
OF: open fieled test
ORF: Open reading frame
PBS: phosphate bufferedsSaline
PCI: peritoneal contamination and infection
PFA: paraformaldehyde
qPCR: quantitative real-time PCR
RIS: RNA Integrity Score
rsn: robust spline normalization SAE Sepsis-associated encephalopathy
SC: Schaffer collateral
s.c.: subcutaneous
SEM: standard error of the mean
TNFα: tumor necrosis factor-α
trkB: Tropomyosin receptor kinase
TTX: Tetrodotoxin
WPRE: Woodchuck Hepatitis Virus Posttranscriptional Regulatory Element

## Acknowledgements

We thank C. Sommer, C. Reißig and D. Himsel for providing expert technical assistance.

## Supplementary material

Supplementary material is available at Brain online.

## Supplementary Material

### Supplementary Material and Methods

#### Behavioral analysis

##### Open Field (OF)

OF testing was performed similar to a previously described protocol.^1^ Here, a custom-made square-shaped box (50×50 cm) was used with walls impermeable for light. The box was open to allow filming and illumination. The arena was divided into a center zone (25×25 cm) and the surrounding periphery zone. Due to shadows arising from the walls of the OF, illumination at the floor level slightly decreased from the center (180-200 lux) to the periphery (120-150 lux). Individual animals were placed in the arena adjacent and oriented towards the wall. Mice were allowed to explore the OF for 5 minutes while the animals were videotaped and tracked (EthoVision, Noldus, Wageningen, Netherlands, or ANYmaze, Stoelting Europe, Dublin, Ireland). Parameters measured included total time spent in center zone or periphery zone, total distance travelled, and number of entries into the zones. After completion of a trial, OF was cleaned with 70% ethanol to ensure identical experimental conditions for the subsequent trial.

##### Elevated Plus Maze (EPM)

A custom-made EPM consisting of grey plastic was used. The EPM was elevated 100 cm above the floor level and had two opposing open and two closed arms (31 x 5,8 cm) as well as an intersection (5,8 x 5,8 cm). The closed and open arms, and the intersection were illuminated with 40, 160 and 120 Lux, respectively. Mice were placed in the intersection facing to an open arm and locomotion was tracked for 5 minutes. The number of arm entries and the time spent in closed and open arms as well as the intersection were recorded using a camera tracking system (EthoVision or ANYmaze). An arm entry was defined by all four feet being set in the respective arm. The EPM was cleaned using 70% ethanol after each trail to ensure equal experimental conditions.

##### Morris Water Maze (MWM)

A custom-made MWM of 119 cm diameter was used which was covered with white foil and illuminated with 30 Lux. The water temperature was set to 20 °C and water was supplemented with milk for better tracking of C57Bl/6J mice and to prevent that mice can visually identify the hidden platform. The diameter of the platform was 10.5 cm and it was placed 1.5 cm below the water surface. Four quadrants were subdivided by clearly distinguishable landmarks placed around the MWM. On day −1 mice were placed in the MWM for 60 s without hidden platform for habituation. Thereafter, mice were subjected to testing in the MWM for 7 consecutive days with 3 trials per day. Here, mice were placed in the water facing towards another quadrant in every trial in a predetermined manner. Time and the distance moved until mice reached the platform were determined. Mice had to enter the platform and had to stay on the platform for 3 s for evaluation of the trial being successful. Trial was terminated by the investigator when mice did not find the platform within 60 s and mice where then gently moved to the platform to assure learning in every trial. On the following day (day 8) a probe trial was conducted. Here, the hidden platform was removed and the time spent and distance travelled in the target quadrant (previous position of the platform) was determined. Tracking was performed using ANYmaze.

##### Barnes Maze (BM)

A circular white PVC platform (diameter 120 cm), mounted on a rotatable stand, elevated 100 cm above floor level, and consistently illuminated by 2 light sources (900-1000 Lux) was used. Three different cues (black-colored geometric shapes printed on paper) were placed at three walls surrounding the maze. At the open side of the maze the examiner was regarded as the cue. The maze comprised 40 pseudo-randomly distributed and equally sized holes (diameter each 5 cm). During the habituation (d0) and training phase (d1-6) 3 an escape box was mounted below one of the holes, while the other holes were closed with plastic disks made of the same material (black PVC). 4 different holes at the respective sides of the maze were used as escape holes. The location of the escape box was kept constant for each mouse but varied between mice (total of 4 possible target locations). Before starting each experiment, mice were acclimated to the testing room for at least 30 minutes and each mouse was placed in an individual holding cage after the experiment. After each trial the maze was cleaned with 70% ethanol to ensure comparable experimental conditions. In the habituation phase (d0), mice were placed in a transparent plastic located in the center of the maze. After 60 seconds the cylinder was lifted and mice were allowed to explore the maze for 60 seconds and then gently led to the escape box using the transparent cylinder. Mice were allowed to spend 2 minutes in the escape box before they were placed in individual holding cages. Each mouse underwent the habituation phase thrice with an inter-trial interval of approximately 10 minutes. In the training phase (d1-6), an opaque instead of the transparent plastic cylinder was used. The cylinder was lifted after 20 seconds and mice were videotaped and tracked (EthoVision) for 4 minutes or until the mouse entered the escape box. In the testing phase (d7) all holes were closed. After 20 seconds in the opaque cylinder, mice were videotaped for 60 sec and the time spent and distance travelled in the target quadrant (previous position of the escape box) was determined.

### Immunohistochemistry

The brains were post-fixed in 4% PFA for 24h at 4°C, cryoprotected in 10% and 30% sucrose in PBS and stored at −80°C. Coronal 40 µm slices were prepared on a sledge microtome equipped with a freezing unit (Thermo Fisher Scientific, Kalamazoo, USA). Slices were incubated in blocking buffer (TBS, 0,1% Triton X, 3% Serum, 2% skimed milk powder) for 30 min. First antibody was incubated in blocking buffer over night at 4°C (anti-ARC/Arg3.1, Synaptic System, Göttingen, Germany, 1:500). Slices were washed in TBS and incubated with 2^nd^ antibody for 2h at room temperature (Rhodamin@ guinea pig, Jackson ImmunoResearch, 1:300). Slices were mounted on gelatin-coated glass slides, stained with DAPI and embedded using Mowiol.

### Golgi staining and dendritic spine analysis of CA1 pyramidal neurons

Morphological spine analysis was performed in pyramidal neurons of the CA1 region as described previously.^2^ Directly after removal of the mouse brain, brains were cut into hemispheres and one hemisphere was used for Golgi staining. Golgi silver impregnation was conducted with the FD Rapid GolgiStainTM Kit (FD NeuroTechnologies, Inc., Columbia, USA) according to manufacturer’s instructions. Briefly, the tissue was rinsed in ACSF to remove blood residues, and was then transferred into equal amounts of solution A+B of the FD Rapid GolgiStainTM Kit. After renewal of solutions on the second day the tissue was subsequently left in the dark for 2 weeks at room temperature. Thereafter, brain hemispheres were transferred into solution C of the FD Rapid GolgiStainTM Kit, which was changed after one day, and were left in solution at C for 5 days. Hemispheres were then stored at 80°C until further use. Then, hemispheres were cut into 150mm thick coronal slices in the area of the hippocampus using a Leica CM3050S cryostat and mounted on object slides. For the last step of staining procedure, slices were washed in purified water (2×4 min), incubated in a mix of 1/4 solution D, 1/4 solution E and 1/2 purified water for 10 minutes, washed again, and then dehydrated in an ascending alcohol series (50%, 75%, 95% for 4 min each, and 100% for 8 min). Finally, slices were immersed in xylene (2×5 min), and then quickly coverslipped with Entellan^®^ (Merck KGaG,Darmstadt, Germany) and dehumidified overnight. To evaluate dendritic morphology, a Zeiss Axioskop 2 mot plus and a computer-based system (Neurolucida; MicroBrightField) was used to generate three-dimensional neuron tracings that were subsequently analyzed using the NeuroExplorer Software (MicroBrightField). Golgi-impregnated cells were selected for reconstruction if they fulfilled the following criteria: (1) the neuron was located within the pyramidal cell layer of the CA1 region (2) the neuron was distinguishable from neighboring cells to allow for identification of dendrites, (3) the dendrites were not truncated or broken, and (4) the cell exhibited dark and well filled impregnation throughout whole dendrites including spines. For analysis of dendritic spines, positively stained neurons in the CA region were randomly selected. Neuronal apical dendrites were followed up to tertiary branches without visible disruptions and without further split up using a Zeiss Axioskop 2 mot plus with an 100x oil immersion objective (PLAN-Neofluar, Zeiss). 3^rd^ order branches of 20-50 µm stretch, 50-150 µm away from the soma of putative CA1 pyramidal cells were analyzed using Neurolicida Neuron Tracing Software. Spines were classified as follows: 1. Thin: shorter than 1,5 without swellings, 2: Mushroom: spine head larger than 0,6 µm 3. Stubby: length/width smaller than 1. Number and type of spines (mushroom, stubby, thin) were counted manually by the same blinded investigator. Total spine density was calculated as number of spines per 10 mm of dendritic length.

### Western Blot

Brains were snap frozen in liquid nitrogen and stored at −80°C until further use. Lysates were obtained using ice-cold RIPA buffer supplemented with complete protease inhibitor Kit and PhosSTOP inhibitor cocktail (both: Roche Diagnostics, Mannheim, Germany). The protein concentration was determined by Bradford Assay (BioRad Quick Start Bradford 1x Dye Reagent, #5000205). Proteins samples (30µg per lane) were subjected to SDS-Page and transferred to a PVDF membrane (Amersham™ Hybond^®^, GE Healthcare Life Science, United Kingdom). First antibodies were incubated overnight at 4°C (ARC/ARG3.1, 1:7500, #156005; Synaptic Systems GmbH, Germany; GAPDH, 1:10000, #2118L, Cell Signalling Technology, USA). Horse-radish peroxidase-labelled secondary antibodies (1:1000, Dianova, Germany) were incubated for 2h at room temperature. Chemiluminescent signals (Pierce ECL Western Bloting Substrate, Thermo Scientific, USA) were visualized using a luminescent image analyzer (LAS 3000, Fujifilm Life Science). Densitometric analysis was performed using ImageJ software. Regarding ARC expression 10 weeks after the PCI the summed signal of the ARC double band was used for analysis.

### Capillary western immunoassay

Mouse hippocampi were homogenized in lysis buffer (0.32 M Sucrose, 4 mM Tris-HCL (pH = 7.4), 1 mM EDTA, 0.25 mM DTT) with additional ultrasound sonication (5x pulses). The protein concentration was determined using Bradford assay (BioRad Quick Start Bradford 1x Dye Reagent, #5000205) prior to protein analysis by capillary western immunoassay (Wes^TM^, ProteinSimple) according to the user manual. For each target, the antibody dilution and dynamic range has been determined independently to assure quantitative measurements. Proteins (0.2µg/µl) were separated in the 12-230 kD detection module (ProteinSimple, # SM-W004) and detected in multiplex assays with a-Tubulin (Cell Signaling Technology, #2144S, 1:10), Erk (Cell Signaling Technology, #4696, 1:10), and pErk (CellSignaling, #4370, 1:15) antibodies, respectively. The antibody incubation time of primary as well as of secondary antibodies was 30 min. The data from two separate experimental cohorts were normalized to the respective averaged SHAM levels. The normalized data were used for statistical analysis.

## Supplementary figures and legends

**Supplementary Figure. 1.**
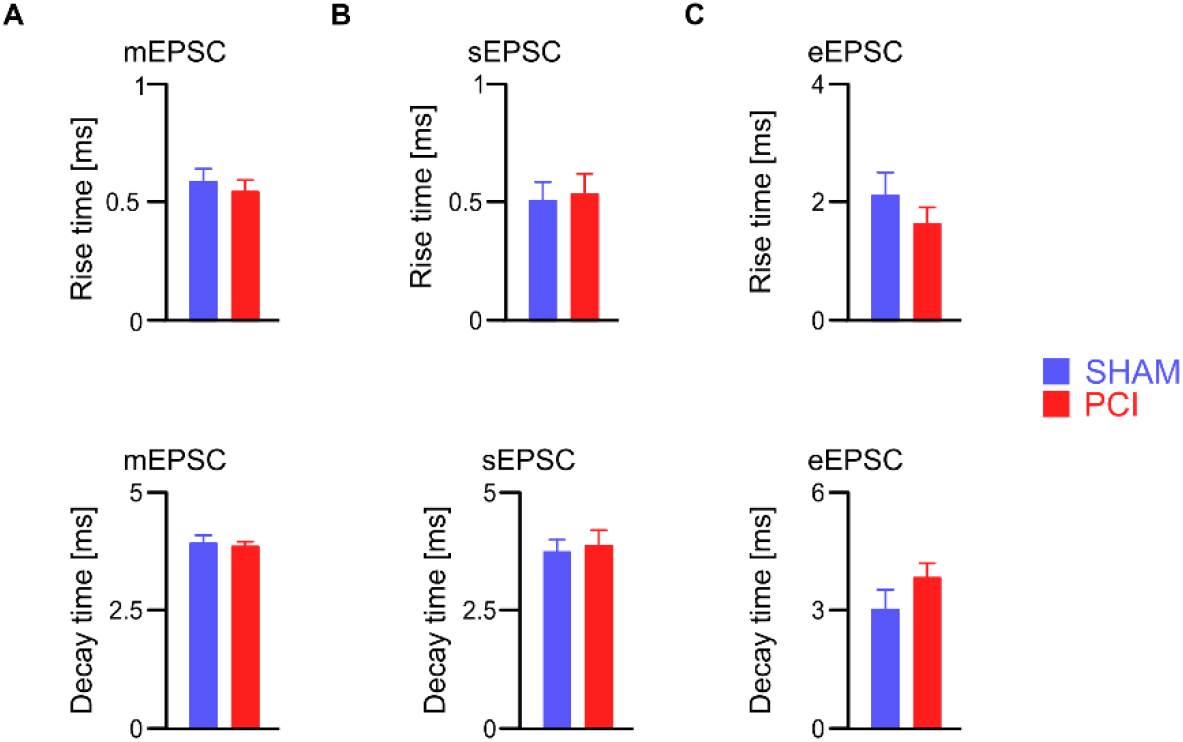
Unchanged time course and kinetics of mEPSC, sEPSC and eEPSC. (**A**) mESPC rise and decay time. (**B**) sEPSC rise and decay time. (**C**) eEPSC rise and decay time (mean ± SEM).

**Supplementary Figure. 2.**
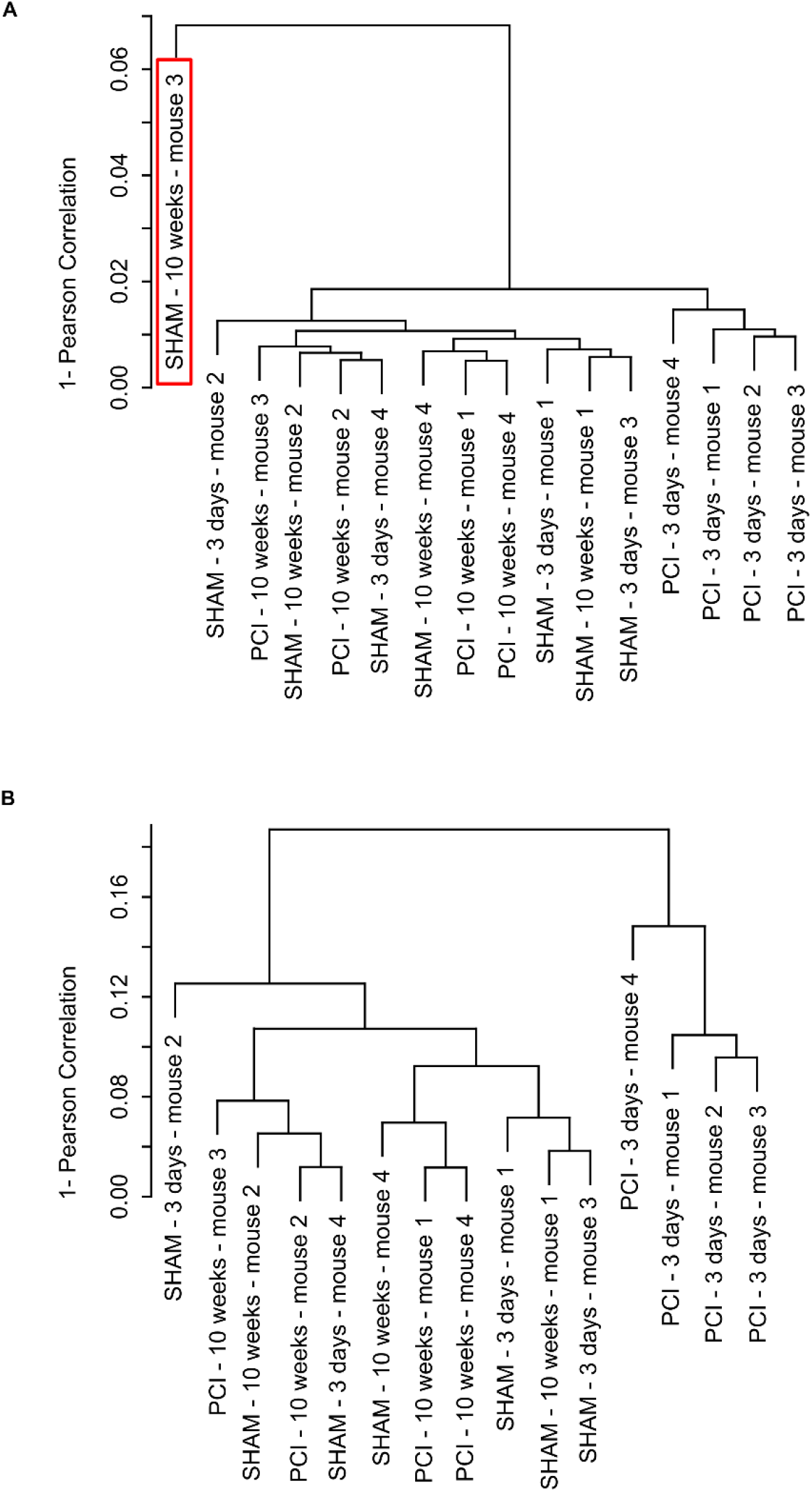
Outlier identification by hierarchical cluster analysis of microarray data. (**A**) Cluster dendrogram with AU/BP values (distance: Pearson’s correlation; cluster method: average linkage) containing data from 4 brain samples per group (PCI 3 days, PCI 10 weeks, SHAM 3 days, SHAM 3 weeks). Clustering identified one data set as an outlier (Sham – 10 weeks mouse 3, Chip-Pos.: 9298740075_G; red border) which was thereupon removed from further analysis. (**B**) Cluster dendrogram without outlier ‘9298740075_G’. All microarray data are available with the following accession number: GSE167610 (NCBI GEO DataSets).

**Supplementary Figure 3.**
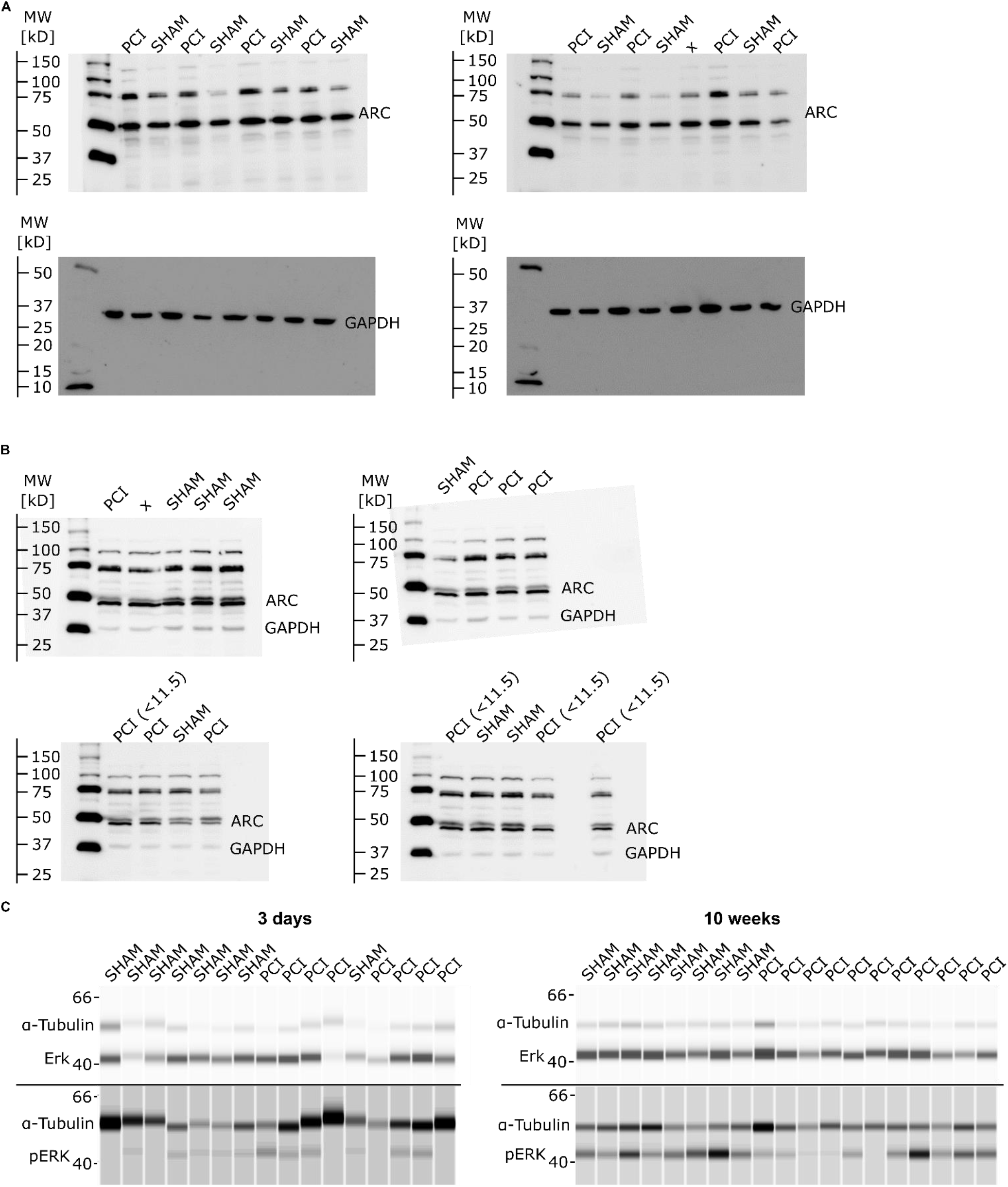
Protein analysis. (**A**) Full Western blot of ARC expression in brain tissue 3 days after PCI. GAPDH was used as loading control and stained for on the same blot after stripping (treatment of groups as indicated). (**B**) Full Western blot of ARC expression after 10 weeks of recovery are shown. GAPDH is stained on the same blot. (Note: only PCI mice with a CSS Score above 11.5 were included). (**C**) Lanes of capillary western immunoassay showing expression of ERK and pERK, as indicated.

**Supplementary Figure 4.**
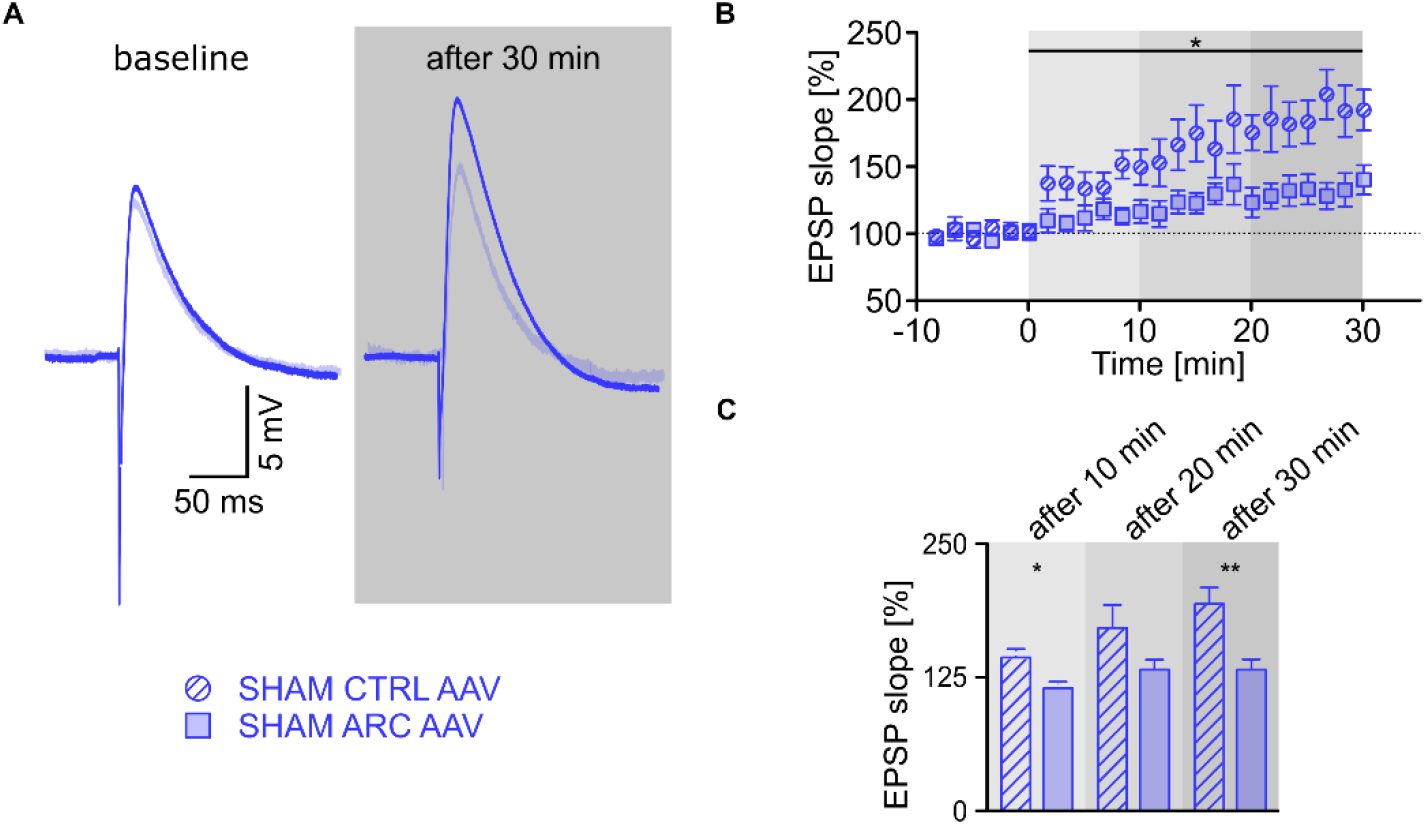
Constitutive hippocampal ARC overexpression impairs LTP in CA1 neurons of non-septic control mice. (**A**) LTP as measured by slope of excitatory postsynaptic potential (EPSP) in single-cell whole cell recordings in CA1 neurons after stimulation of Schaffer collaterals. Average traces of 10 recordings before and 30 min after theta burst stimulation. (**B**) LTP time course. LTP is reduced in SHAM ARC AAV (n = 11) mice in comparison to SHAM CTRL AAV (n = 11; 2-way repeated measurement ANOVA; Tukey’s post hoc test). (**C**) Comparison of LTP at the indicated time points after theta burst stimulation. *P < 0.05, **P < 0.01; 2-way repeated measurement ANOVA; Tukey’s post hoc test.

**Supplementary Figure 5.**
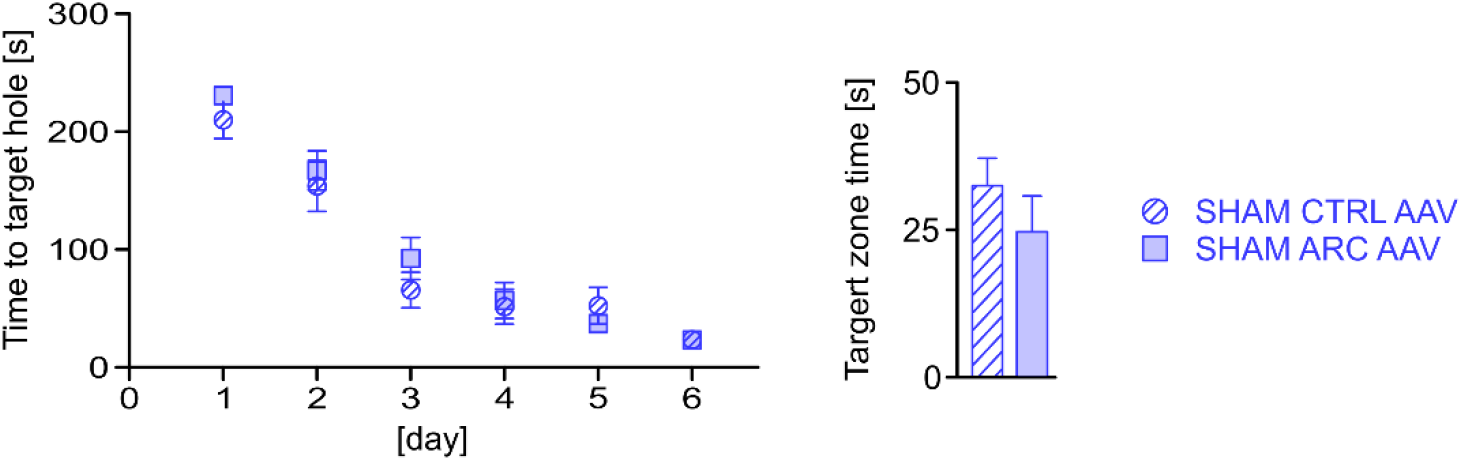
Overexpression of ARC in SHAM mice does not improve learning and memory. Barnes maze (BM) testing of SHAM CTRL AAV (n = 11) and SHAM ARC AAV (n = 11) animals reveals unchanged learning abilities in the training phase (d1-6, left) and memory function in the testing phase (probe trial, d 7, right).

**Supplementary Figure 6.**
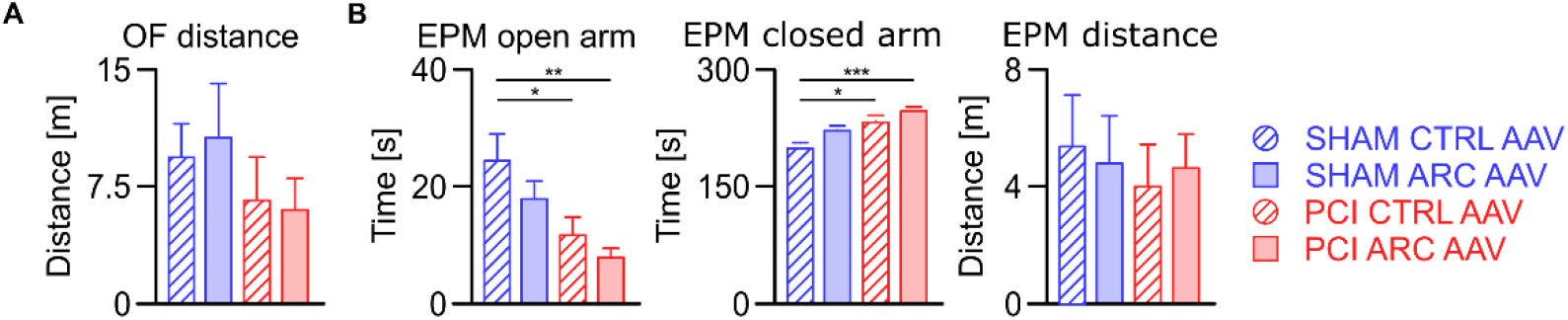
Unchanged locomotor activity and persistent anxiety-related behavior in PCI after hippocampal overexpression of ARC. (**A**) Unchanged locomotor activity in the OF indicating loss of general sickness behavior in all experimental groups (same number of mice in the respective treatment groups as detailed in B). (**B**) PCI mice with control AAV injection (PCI CTRL AAV; n = 13) spend reduced time in the open arms of the elevated plus maze (EPM) indicating fear-related behavior. As expected, sepsis-induced anxiety-related behaviour did not improve upon hippocampal overexpression of ARC (PCI ARC AAV; n = 16). Control mice do not show abnormalities in the EPM (SHAM CTRL AAV, n = 17; SHAM ARC AAV, n = 18). Locomotor activity in the EPM (EPM distance) is unchanged in all groups. * P < 0.05, 2-way ANOVA with Sidak post hoc test.

**Supplementary Figure 7.**
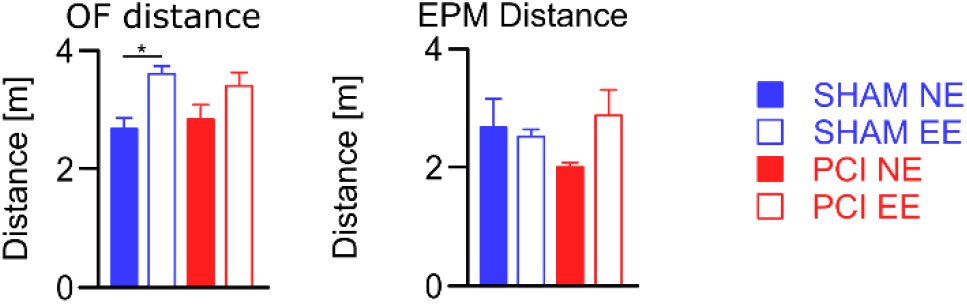
Unchanged locomotor activity in PCI animals after enriched environment compared to SHAM animals. Unchanged locomotor activity in the OF and EPM upon PCI indicating loss of general sickness behaviour in all experimental groups. Note, that enriched environment leads to general slight increase of mobility which is statistically significant in the SHAM EE group in the OF testing (SHAM NE, n = 15; SHAM EE, n = 10; PCI NE: 14; PCI EE, n = 18); *P < 0.05, 2-way ANOVA with Tukey’s post hoc test.

